# MORC2 directs transcription-dependent CpG methylation of human LINE-1 transposons in early neurodevelopment

**DOI:** 10.1101/2025.06.11.658943

**Authors:** Fereshteh Dorazehi, Caterina Francesconi, Ninoslav Pandiloski, Laura Castilla-Vallmanya, Anna Albecka, Symela Koutounidou, Sivani Bala Mohan, Anita Adami, Angelos Katsikas, Jana Matijević, Ofelia Karlsson, Carrie Davis-Hansson, Jenny G. Johansson, Yorgo Modis, Johan Jakobsson, Christopher H. Douse

**Author notes:** these authors contributed equally (Caterina Francesconi, Ninoslav Pandiloski, Laura Castilla-Vallmanya).

## Abstract

Methylation of CpG dinucleotides is essential for silencing genomic repeats such as LINE-1 retrotransposons (L1s) in the germline and soma. Evolutionarily-young L1s are transcribed in human pluripotent stem cells, but how CpG methylation is patterned to these L1s upon exit of pluripotency is unknown. Here we investigate the critical functions of chromatin regulator MORC2 in epigenome reprogramming of the repetitive genome in early human neurodevelopment. We find that reversible ATP-dependent dimerization is required for MORC2 accumulation over L1s but not gene promoters. Engineered mutations of the MORC2 ATPase module severely disrupts the distribution of MORC2 chromatin binding, leading to simultaneous loss of L1 transcriptional control and hyper-repression of clustered ZNF genes in human pluripotent stem cells. Upon neural differentiation these phenotypes persist due to striking, targeted defects in CpG methylation patterning. Together our results define the vital role of MORC2 in safeguarding the somatic human genome upon exit of pluripotency by directing CpG methylation patterning over transcriptionally-active retrotransposons in a manner analogous to the piRNA pathway in the germline.

## INTRODUCTION

Transposable elements (TEs) – in particular, retrotransposons such as long interspersed nuclear element 1 (L1) – have accumulated in mammalian genomes and account for at least half of human DNA.^1^ Out of more than half a million fixed L1s in the reference genome, around two thousand hominoid-specific copies retain an intact internal promoter and are thus competent for transcription. These elements are a source of novel transcripts in human tissues.^2–5^ A smaller subset of around 100 human-specific L1s encode proteins that enable transposition of their own and other TE transcripts to new sites, resulting in polymorphisms within and between individuals.^6,7^ L1 fragments lacking their own promoter often reside in introns of transcribed genes where they can influence mRNA processing and gene expression.^8–11^ L1 activity is therefore responsible for substantial genetic and transcriptional variation in the human population.

Unrestricted mobilization of L1s and other TEs is thought to be deleterious and transcription of intact sequences is controlled by promoter CpG methylation in somatic mammalian cells.^12–15^ As part of the epigenetic reprogramming that happens during germ cell and embryonic development, when TE transcription is transiently tolerated and in certain cases functionally important,^16–20^ DNA methylation patterns are reestablished after periods of global erasure. In the germline, the piRNA pathway licenses transcription-dependent TE methylation.^21^ In embryonic stem cells, the most studied mechanism of restriction works via the KRAB zinc-finger (KRAB-ZNF) family of transcription factors, which bind to specific TE DNA sequences and recruit the TRIM28 corepressor that promotes histone 3 lysine 9 trimethylation (H3K9me3) and CpG methylation.^22,23^ KRAB-ZNF genes have rapidly diversified by segmental duplication during vertebrate evolution to recognize TE and repeat family sequences.^24,25^ However, in the human setting, it is thought the youngest L1 subfamilies have evaded this DNA-directed silencing system: changes in the ancestral L1PA3 promoter allowed subsequent, evolutionarily-younger L1 copies to escape silencing by ZNF93.^26,27^ In human pluripotent stem cells, L1 expression is indeed mostly restricted to human-specific (L1HS) and hominoid-specific (L1PA2/3) elements, which correlates with relative promoter hypomethylation and varies between individual integrants.^28–30^ Although lineage-specific L1-driven transcripts exist, and intronic L1s are passively transcribed in pre-mRNA of thousands of genes, CpG methylation mostly silences the promoters of the youngest, intact L1s in somatic human tissues.^13,31^ How methylation is patterned to young L1 promoters upon exit of pluripotency and differentiation has been an open question, but is not thought to depend solely on the KRAB-ZNF/TRIM28 system.

A key player in the regulatory relationship (sometimes referred to as an ‘arms race’^26^) between L1s and KRAB-ZNF genes is the chromatin remodelling ATPase MORC2.^8,32^ In collaboration with the human silencing hub (HUSH) complex, MORC2 is recruited to repress transcriptionally-active L1s and repetitive gene clusters, including those encoding KRAB-ZNFs, at H3K9me3-marked sites.^8,32–35^ ChIP-seq experiments in cancer cell lines also identified a second class of MORC2 peaks over transcription start sites, the functional significance of which is unknown.^32^ MORC2 contains an N-terminal ATPase module that reversibly dimerizes upon ATP binding and hydrolysis, and a C-terminal coiled coil domain that forms constitutive dimers.^36–38^ Elegant recent experiments with purified components have shown that MORC2 binds and entraps free and nucleosomal DNA between these dimerization domains in a length but not sequence-dependent manner, forming loops and multimeric assemblies that compact the underlying chromatin.^36^ This mechanism of MORC chromatin remodelling appears to be conserved at least across the animal kingdom^39^ and perhaps more deeply – MORCs are encoded in plant, basal eukaryote and even prokaryote genomes.^40–43^

MORC2 functions are vital to human health: predicted loss of function alleles are strongly selected against in the population^44^ and heterozygous missense mutations in the MORC2 ATPase module cause severe neurodevelopmental disorders with symptoms including intellectual disability and cerebellar ataxia.^45–47^ Patient mutations cause changes to the protein’s enzymatic activity and hyperrepression of clustered KRAB-ZNF genes in cancer cell lines.^32^ A recent paper profiling peripheral blood and fibroblasts from MORC2 disorder patients revealed an ‘episignature’ corresponding to hypermethylation and repression of genes including KRAB-ZNFs, showing that MORC2 neurodevelopmental disorders are characterized by striking epigenetic changes in cells outside the nervous system.^47^ Notably, the functional consequences of removing MORC2 depends substantially on the cellular context. For example, loss of MORC2 in the globally-hypomethylated K562 line causes massive upregulation of young L1 subfamilies, together with changes in KRAB-ZNF expression.^8^ In contrast, we recently found that removal of MORC2 in a robustly-methylated neural progenitor cell line had only mild effects on L1 expression – indeed, only when DNA methylation was experimentally removed did MORC2 bind to the activated L1s and dampen their expression.^33^ Thus, when young L1 promoters are methylated and silenced, MORC2 ignores these sequences and is bound to either older, intronic L1s (transcribed as part of genic pre-mRNA) or other transcribed lineage-specific repeats – in the nervous system, data from our lab and others suggest these to be primarily alpha-satellite repeats and clustered protocadherin genes.^33,34^

Together these observations underscore the layered regulation of repeats in somatic cells and suggest critical functions of MORC2 early in development when the genome is relatively hypomethylated. In this study we use a combination of chromatin profiling, long-read DNA methylation analysis and iPSC modelling to demonstrate (i) how ATPase mutations affect the distribution of MORC2 chromatin binding in the neural lineage, and (ii) the critical function of MORC2 chromatin binding in directing CpG methylation patterns over hominoid-specific L1s and repetitive gene clusters in early neurodevelopment. Together our results define the vital role of MORC2 in safeguarding the somatic human genome upon exit of pluripotency by ensuring CpG methylation patterning over repeats in a transcription-dependent manner, analogous to the piRNA pathway in the germline.

## RESULTS

### MORC2 binds to distinct chromatin states in neural cell lines

MORC2 is a potent and avid dsDNA binding protein.^36,37^ The prevailing biochemical model is that reversible ATP-dependent dimerization of the N-terminal ATPase module coupled to constitutive C-terminal dimerization via coiled coil domain 3 enables chromatin compaction by trapping DNA between these domains (**Figure 1A)**.^36,39^ We first profiled MORC2 chromatin binding using an optimized CUT&RUN (cleavage under targets and release using nuclease) protocol in SHSY5Y cells and embryo-derived human neural progenitor cells (**Figure 1B, Supp Fig 1A,B**).^48^ In each cell type we clustered MORC2 binding regions by assessing peak shape and co-occupancy with three histone modifications in control cells: H3K9me3, H3K4me3 and H3K27Ac (**Figure 1C**). We observed broad MORC2 peaks in transcriptionally active, H3K9me3-positive regions such as intronic full-length (>6 kb) L1 retrotransposons and 3’ exons of expressed zinc finger (ZNF) genes (**Figure 1D, Supp Fig 1C)**. We also observed widespread evidence of MORC2 binding to bivalent chromatin i.e. H3K4me3/H3K27Ac-marked promoters of expressed genes embedded in H3K9me3-positive chromatin, such as clustered ZNF and protocadherin genes. In our hands a minority of narrow MORC2 peaks were identified over gene regulatory elements marked by H3K27Ac and/or H3K4me3 in H3K9me3-negative chromatin contexts. Taken together, in neural cells MORC2 localizes to distinct chromatin states characterized by transcriptional activity and either broad or narrow occupancy, exemplified by L1 retrotransposons and repetitive clustered genes such as ZNFs.

**Figure 1.**
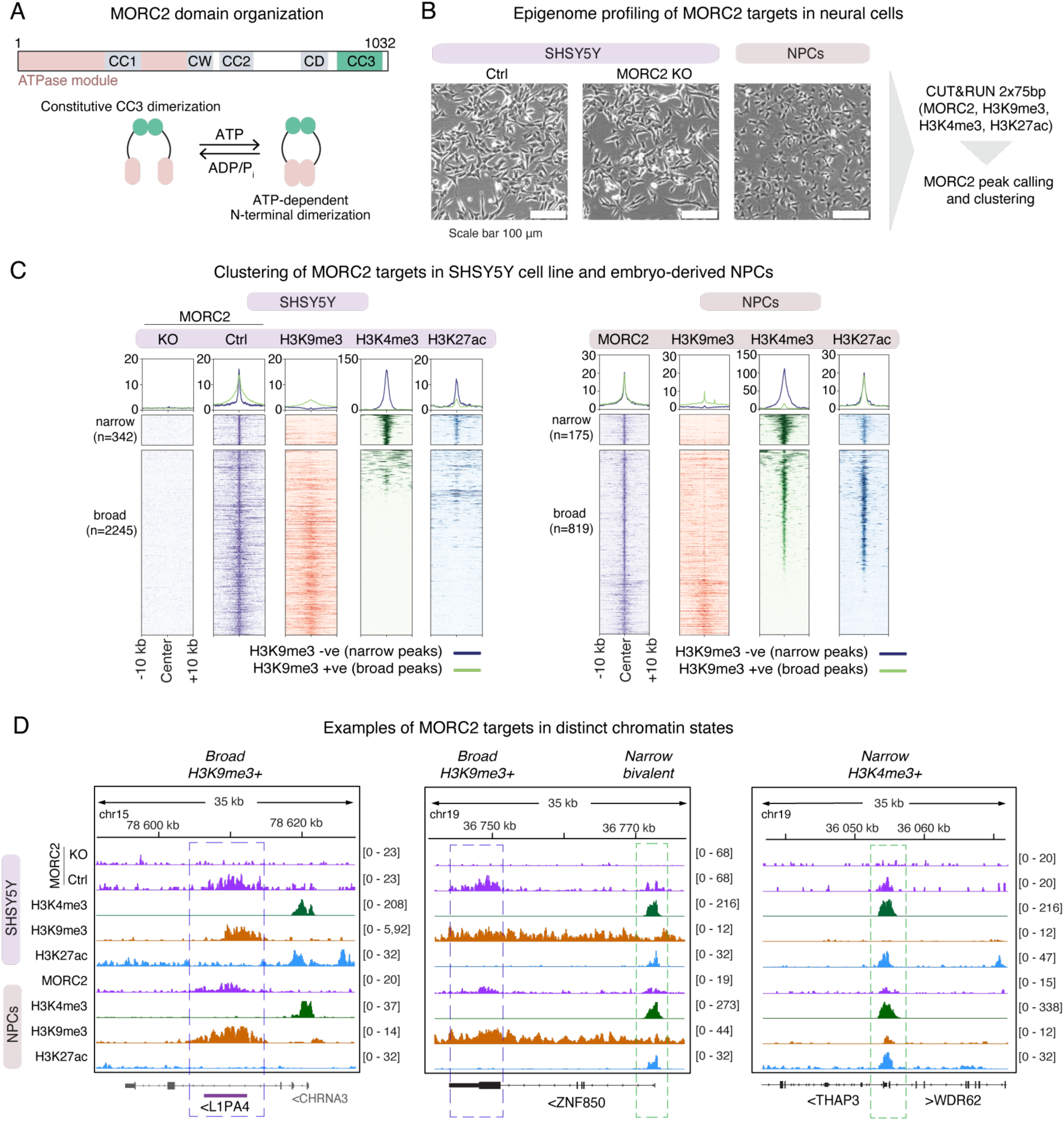
MORC2 binds to distinct chromatin states in neural cell lines. **(A)** MORC2 domain structure (top) and schematic of ATP-dependent N-terminal dimerization (bottom). **(B)** CUT&RUN workflow for epigenome profiling of SHSY5Y and hNPC cell lines. (**C)** Heatmap illustrating the epigenomic context of distinct MORC2 targets by CUT&RUN signal in SHSY5Y (left) and NPC (right) cell lines with indicated antibodies. Displayed are the genomic regions spanning ± 10 kb from the center. K-means clustering defined H3K9me3-negative, MORC2-narrow peaks (top) and H3K9me3-positive, MORC2-broad peaks (bottom) in each cell type. **(D)** Genome browser tracks showing CUT&RUN enrichment profiles of MORC2 and histone modification over examples MORC2 targets with distinct epigenomic associations. Data are representative of at least two replicates for MORC2, H3K9me3 and H3K4me3 in each line, while H3K27Ac profiling was conducted once in SHSY5Y cells and twice in hNPCs.

### ATPase dimerization mutants abolish L1 binding and disrupt the balance of MORC2 chromatin occupancy

Mutations associated with MORC2 neurodevelopmental disorders all cluster to the N-terminal ATPase module.^46^ To interrogate how reversible ATP-dependent dimerization impacts MORC2 chromatin binding we made use of two mutants that we previously found shift this equilibrium in opposite directions: Y18A, a structure-guided mutation that prevents ATPase dimer formation, and S87L, a patient mutation that leads to constitutive ATPase dimerization (**Figure 2A**).^37^ We reintroduced HA-tagged proteins to clonal MORC2-KO SHSY5Y cells (**Figure 2B**) and first confirmed that WT, Y18A and S87L MORC2 variants were expressed and nuclear as assessed by Western blotting (**Figure 2C**) and imaging (**Supp Fig 2**). CUT&RUN analysis revealed that both Y18A and S87L variants were completely lost over L1s bound by the WT protein (**Figure 2D,E**). Importantly, these effects were reproducible using antibodies raised against the MORC2 protein itself and its HA epitope tag. Both variants were lost from 3’ exons of ZNF genes but remained bound to the promoters of those genes, suggesting that ATPase dimerization is not required for promoter occupancy. Notably, the binding of S87L MORC2 to ZNF promoters was reproducibly enhanced relative to WT, suggesting that this mutation causes relocalization of the protein away from L1s and onto gene promoters (**Figure 2F,G**). Indeed, peak calling revealed a substantially greater overlap of S87L MORC2 with promoters when compared to the WT distribution (**Figure 2H**). Taken together, these results demonstrate that ATPase dimerization mutants disrupt the balance of MORC2 chromatin occupancy. Reversible ATP-dependent dimerization is required for broad MORC2 accumulation over L1s but not promoters of target genes (**Figure 2I**). The S87L mutation directs the protein away from L1s and other broad MORC2 targets and increases binding over gene promoters.

**Figure 2.**
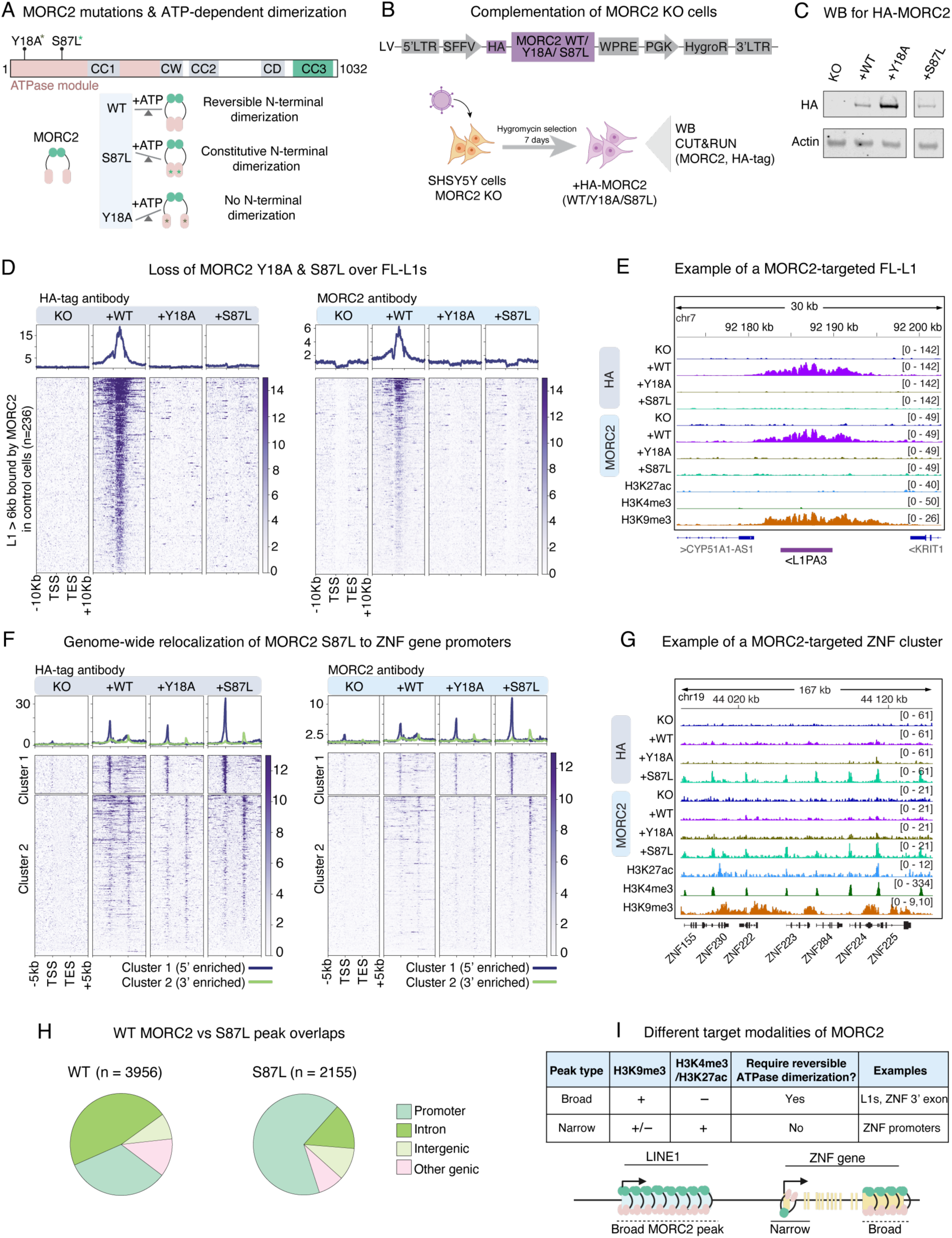
ATPase dimerization mutants abolish L1 binding and disrupt the balance of MORC2 chromatin occupancy. **(A)** Schematic representation of MORC2 protein domains and the position of ATPase disease mutations (top) and the effect of the mutations on the N-terminal dimerization equilibrium (bottom). **(B)** Schematic workflow of MORC2 complementation experiments and subsequent analyses. **(C)** Western blot validating the expression of complemented HA-tagged MORC2 variants. **(D)** Heatmap showing CUT&RUN enrichment of MORC2 binding over full length (FL, >6 kb) LINE1 elements bound by MORC2 WT in KO and complemented cell lines (+WT MORC2, +Y18A MORC2, +S87L MORC2) as detected using HA-tag antibody (left) and MORC2 antibody (right). Shown are the LINE1 loci scaled to 6 kb with 10 kb up-and down-stream of the elements. Unique mapping was enforced and experiments were repeated at least once with similar results. **(E)** Genome browser tracks of CUT&RUN enrichment signal for MORC2 binding (KO and complemented cell lines) and histone modifications (H3K27ac, H3K4me3, H3K9me3) over an example LINE1 element. Unique mapping was enforced. **(F)** Heatmap showing CUT&RUN enrichment of MORC2 binding over all ZNF genes (n=503, hg38) in KO and complemented cell lines (+WT MORC2, +Y18A MORC2, +S87L MORC2) as detected using HA-tag antibody (left) and MORC2 antibody (right). Shown are the scaled ZNF genes ± 5 kb. K-means clustering on the S87L MORC2 sample identified 5’-enriched (top) and 3’-enriched (bottom) ZNF genes. **(G)** Genome browser tracks of CUT&RUN enrichment signal for MORC2 binding (KO and complemented cell lines) and histone modifications (H3K27ac, H3K4me3, H3K9me3) over a ZNF gene cluster on chr19. **(H)** Genome overlaps of WT and S87L HA-MORC2 peaks. **(I)** Schematic of the requirements for reversible N-terminal dimerization over different target classes of MORC2. MORC2 accumulation over broad peaks is found over regions enriched in H3K9me3 signal, such as LINE1s and ZNF 3’exons, and requires reversible N-terminal dimerization. Narrow MORC2 binding to gene regulatory elements such as ZNF gene promoters is found in regions enriched in H3K4me3 and H3K27ac signal, and does not require reversible N-terminal dimerization.

### MORC2 ATPase mutations cause loss of L1 silencing and ZNF hyperrepression in iPSCs

Next we sought to explore the functional impact of chromatin binding on transcriptional regulation in a relevant developmental cell model. We used CRISPR editing to derive induced pluripotent stem cell (iPSC) clones bearing the S87L mutation described above and a second ATPase mutation, E27K, which also causes a severe neurodevelopmental phenotype and would be predicted to similarly disrupt ATPase activity based on its structural position at the N-terminal ATP-induced dimerization interface.^37,46^ We note here that though patients carry heterozygous mutations, our study sought to provide mechanistic insight into how ATPase mutations affect MORC2 function over a developmental timecourse and as such we selected homozygous *MORC2* edited clones (**Figure 3A, Supp Fig 3A**). We subjected unedited controls to the same clonal expansion procedures and confirmed that each iPSC clone expressed pluripotency markers and that the copy number and genomic integrity at predicted off-targets was not affected by the editing (**Figure 3B, Supp Fig 3B,C**). Transcriptomic analysis using paired-end (150x150-bp) polyA+ RNA sequencing (RNA-seq) showed that the ATPase mutations both cause significant derepression of evolutionarily-young, hominoid-specific L1 subfamilies, with the S87L mutation having a more pronounced effect (**Figure 3C, D**). CUT&RUN profiling of the H3K4me3 mark mapped to individual full-length L1 elements provided orthogonal validation of young L1 expression **(Figure 3E)**. In line with chromatin binding data above illustrating complete loss of L1 binding, the S87L mutation caused L1 transcriptional activation at comparable levels to a loss of function control in which we used a validated CRISPRi vector ^33^ to robustly knock down MORC2 expression in iPSCs (**Figure 3E, Supp. Fig 3D-G**). L1 derepression was further confirmed in Western blots and immunofluorescence experiments as measured by L1-encoded protein orf1p in the mutant and CRISPRi conditions relative to controls (**Figure 3F, Supp Fig 3H**). We also observed pronounced derepression of L1 fusion genes where evolutionarily-young L1s act as alternative promoters (**Supp Fig 4A**). Several of these genes have documented or predicted functions in the brain. For example, *GABRR1*, whose promoter is shared with a L1PA2 element, encodes a key factor involved in GABAergic neurotransmission in the central nervous system^49^ and *DYNC1I1*, which contains an intronic L1PA3, encodes a neuron-enriched dynein intermediate chain (**Supp Fig 4B**). Notably, and in line with the chromatin binding data, alongside L1 loss of function we observed simultaneous hyperrepression of zinc finger genes in E27K (n=22), S87L (n=11) or both (n=19) MORC2 mutants (log2 fold-change < -2, padj < 0.05, DESeq2) (**Figure 3G,H**). This effect was a gain of function associated with the S87L and E27K mutants that was not observed in the MORC2-CRISPRi cells. Taken together these data show how MORC2 ATPase mutations change the dynamics of chromatin target engagement, leading to both loss and gain of function effects on transcriptional control of its distinct targets.

**Figure 3.**
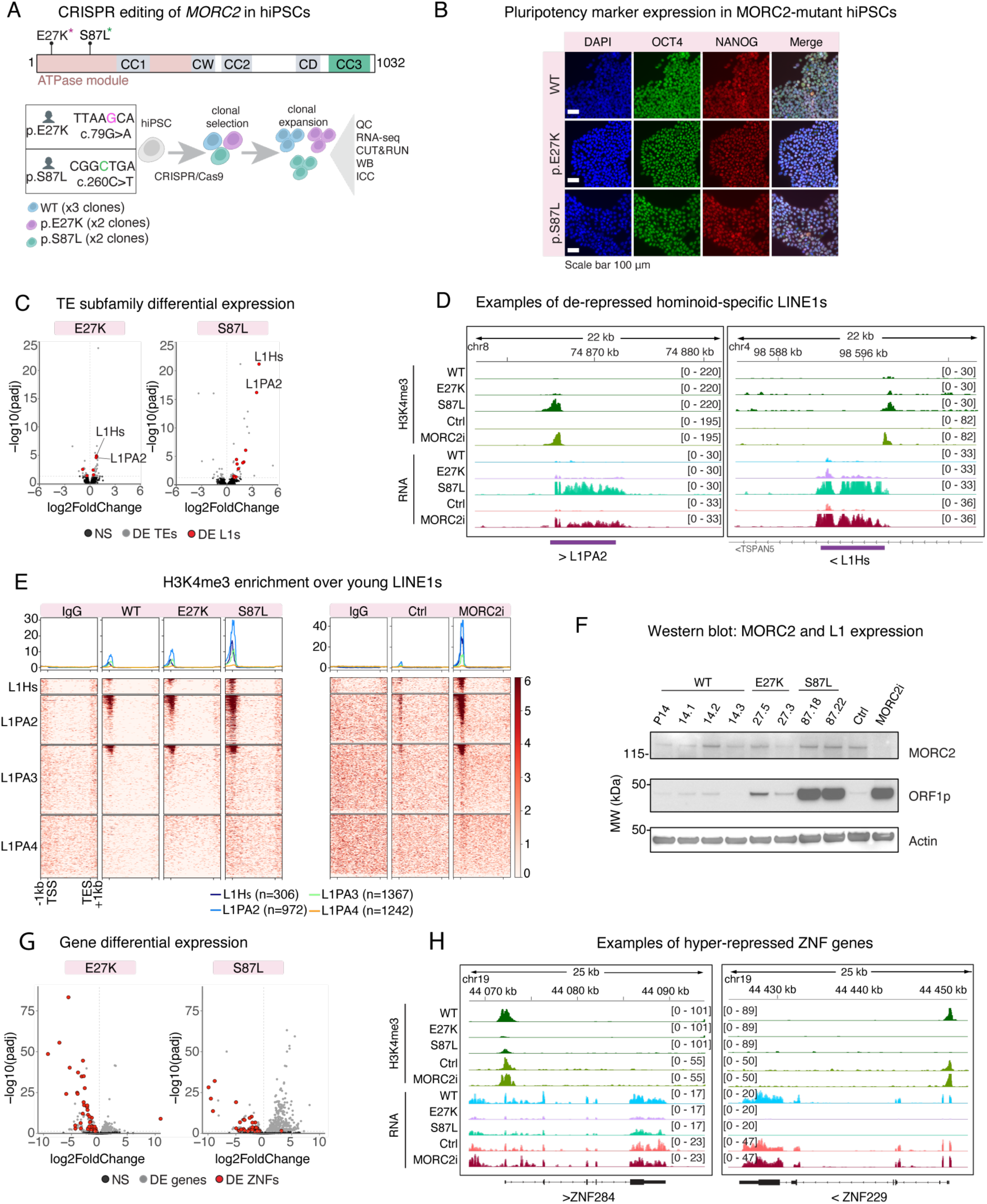
MORC2 ATPase mutations cause loss of L1 silencing and ZNF hyperrepression in iPSCs. **(A)** Schematic of CRISPR editing and subsequent analyses. Induced homozygous mutations were c79G>A and c260C>T for E27K and S87L mutants, respectively. **(B)** Immunocytochemestry of pluripotency markers (NANOG, OCT4, SOX2) for a representative hiPSC clone of each MORC2 genotype. **(C)** Volcano plot showing the results of the differential expression analysis (DESeq2) on a TE subfamily level using TEtranscripts. Differentially expressed LINE1 subfamilies are highlighted in red, other differentially expressed subfamilies are presented in grey (log2FoldChange ≠0, padj <0.05). Two experimental replicates per clone were sequenced and data from each clone pooled prior to differential analysis (n=4 WT vs n=4 E27K vs n=4 S87L). **(D)** Genome browser tracks showing H3K4me3 and bulk RNA-seq data over two example loci of derepressed hominoid-specific full-length L1s. Unique mapping was enforced for all data over individual loci. Also shown are representative data from MORC2 CRISPRi (n=3) and Control (n=3) hiPSCs, see also Supp Fig 3. **(E)** Heatmap of the CUT&RUN signal enrichment of H3K4me3 and non-targeting IgG controls over reference L1 insertions belonging to the evolutionarily-youngest subfamilies (L1HS, L1PA2-4) in MORC2 mutant and CRISPRi cell lines. For each WT and mutant cell line, cells from two clones were pooled prior to CUT&RUN, which was conducted at least twice with similar results. Experiments with control and MORC2 CRISPRi cells were repeated twice with similar results. Unique mapping was enforced for CUT&RUN data shown. **(F)** Western Blot showing expression of MORC2 and of L1 ORF1p in MORC2 mutant and CRISPRi cell lines. The experiment was repeated once with similar results. **(G)** Volcano plot showing the results of the differential gene expression analysis (DESeq2). Differentially expressed ZNF genes are highlighted in red, other differentially expressed genes are presented in grey (padj <0.05). Two experimental replicates per clone were sequenced and data from each clone pooled prior to differential analysis (n=4 WT vs n=4 E27K vs n=4 S87L). **(H)** Genome browser tracks showing H3K4me3 and bulk RNA-seq data over example loci of two hyperrepressed ZNF genes. Unique mapping was enforced for all data over individual loci. Also shown are data from MORC2 CRISPRi (n=3) and Control (n=3) hiPSCs, see also Supp Fig 3.

### Transcriptional phenotypes of MORC2 mislocalization persist upon neural differentiation

To model how these transcriptional phenotypes change upon exit from pluripotency and neural differentiation, we set out to differentiate the iPSC lines to neural progenitor cells (NPCs). Using a previously-described 2D NPC differentiation protocol^50^ we observed widespread cell death in the MORC2-CRISPRi and S87L lines after pluripotency exit, precluding further analysis. As an alternative we made use of an undirected 15-day cerebral organoid protocol as a model for human brain development in 3D (**Figure 4A**). At this timepoint our previous data showed that organoids are largely composed of NPCs, enabling detailed analysis of L1 and gene regulation in bulk.^28^ 3D differentiation improved cell survival and cells bearing both MORC2 mutations or carrying the MORC2 CRISPRi lentiviral vector could form characteristic neural rosettes after 15 days, albeit with morphological and size changes (**Figure 4B**). Single nucleus RNA sequencing confirmed that NPCs and early neurons constituted the major cell populations in these preparations across multiple batches and clones (**Supp Fig 4A-D**). Differential expression analysis on day-15 organoids from at least two clones and three experimental batches showed that the transcriptional phenotype observed in hiPSCs – that is, L1 derepression in mutant and CRISPRi lines and ZNF hyperrepression specifically in the mutants – persisted upon differentiation (**Figure 4C-G**). We found that the promoters of young L1 integrants identified to be derepressed in hiPSCs were highly enriched with the H3K4me3 active mark in MORC2-S87L and MORC2-CRISPRi day 15 organoids (**Figure 4F**). We also observed upregulation of lineage-specific clustered protocadherin genes in MORC2 mutant organoids, while this effect was stronger in MORC2-CRISPRi samples (**Supp Fig 5A,B**). Taken together, our data illustrate that the transcriptional phenotypes of MORC2 targets in the pluripotent state persist upon exit from pluripotency and early neurodevelopment.

**Figure 4.**
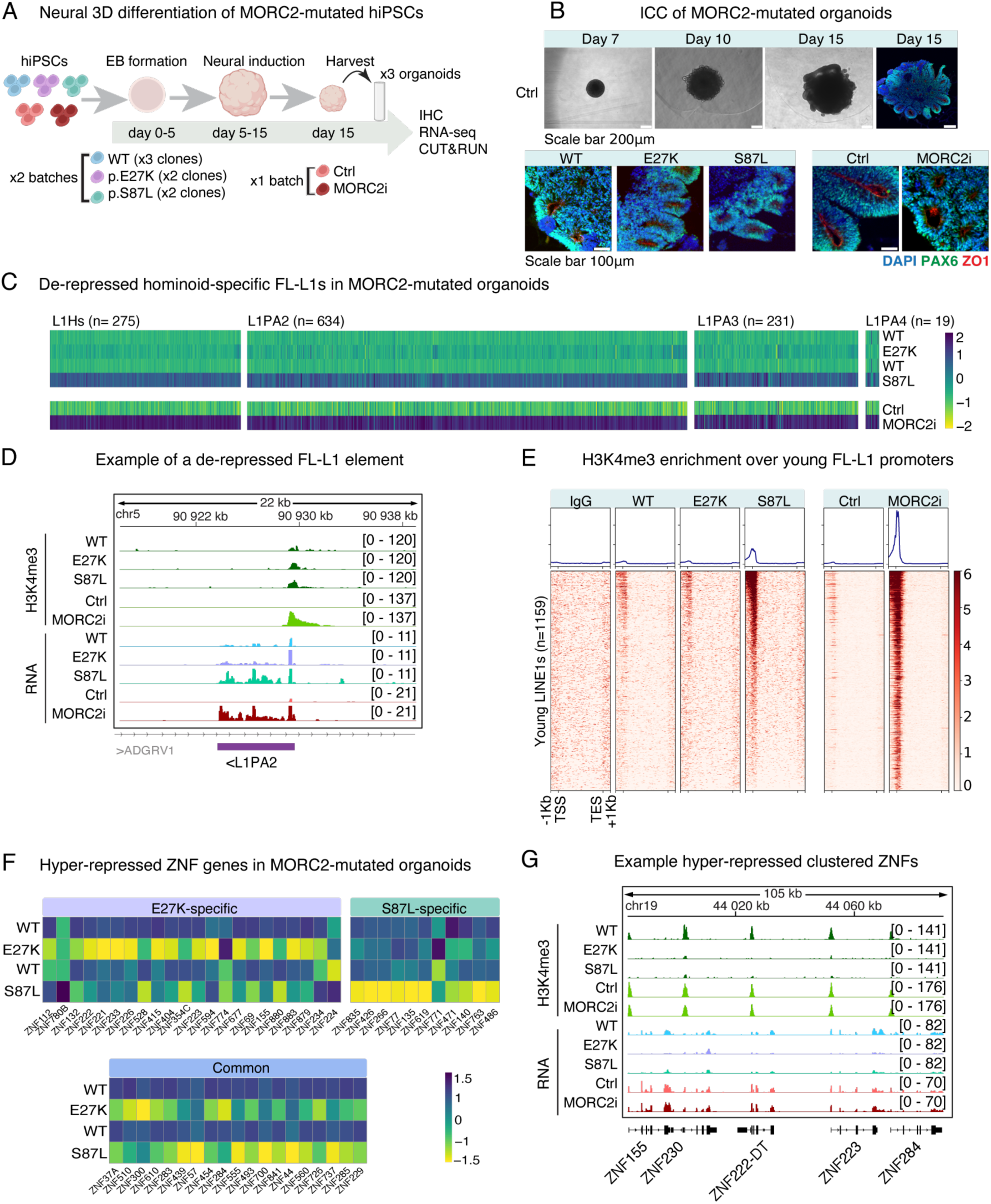
Transcriptional phenotypes of MORC2 mislocalization persist upon neural differentiation. **(A)** Schematic of differentiation of MORC2 mutant and CRISPRi cell lines from hiPSCs to cerebral organoids. Two batches of organoids were produced for both S87L and E27K clones and three for WT clones. One batch was produced for MORC2 CRISPRi and its corresponding control cell line in which hiPSCs were transduced with a non-targeting lentiviral CRISPRi vector. **(B)** Representative bright field images and immunohistochemistry of cerebral organoid formation in a control line at different time points (top). Immunohistochemistry of PAX6 (green) and ZO1 (red) in day 15 organoid sections (bottom). **(C)** Scaled heatmap showing the normalized read counts of individual full length (FL) LINE1 from youngest subfamilies (L1HS, L1PA2-4) that had been derepressed in MORC2 E27K and S87L mutant and MORC2 CRISPRi hiPSCs (log2FoldChange >0, padj <0.05). Data are representative of three replicates of two clone across two batches (n=9 WT vs n=9 E27K vs n=9 S87L). Unique mapping was enforced. **(D)** Genome browser tracks showing CUT&RUN H3K4me3 enrichment and bulk RNA-seq data over an example derepressed full-length L1. Unique mapping was enforced. **(E)** Heatmap of the CUT&RUN signal enrichment of H3K4me3 and non-targeting IgG control over reference full-length L1 insertions belonging to the to the youngest subfamilies (L1HS, L1PA2-4) that were expressed in MORC2 mutants and CRISPRi hiPSCs. For WT and mutant genotypes, organoids from two clones were pooled prior to CUT&RUN, which was conducted at least twice with similar results. Unique mapping was enforced **(F)** Scaled heatmap showing the normalized read counts over ZNF genes hyperrepressed in MORC2 E27K and S87L mutants hiPSCs (log2FoldChange <0, padj <0.05). **(G)** Genome browser tracks showing CUT&RUN H3K4me3 enrichment and bulk RNA-seq data over a hyper-repressed ZNF cluster on chr19. Unique mapping was enforced.

### MORC2 directs CpG methylation patterning over the repetitive human genome

As the observed transcriptional phenotypes persisted from the pluripotent state to day-15 organoids, we hypothesized a defect in CpG methylation patterning over MORC2 targets. To investigate methylation changes over MORC2-targeted repetitive elements we used whole-genome Oxford Nanopore long-read sequencing (**Figure 5A**). While genome-wide methylation was robust and stable across all conditions (**Figure 5B**), the S87L mutation in MORC2 caused a catastrophic methylation loss over the promoter of the youngest L1Hs and L1PA2 integrants (**Figure 5C-E**). Loss of methylation over young L1 promoters was mirrored in the MORC2-CRISPRi organoids and across organoid preparations derived from independent clones, confirming the effect over these targets was reproducible and a direct loss of function effect (**Figure 5C-E, Supp Fig 6A**). Leveraging the Nanopore data to call polymorphic L1 insertions absent from the hg38 reference, we also observed striking loss of methylation over these youngest copies in both genetic backgrounds tested (**Supp Fig 6B**). Importantly, we found that the effect was most striking to the L1 elements transcribed in the pluripotent state, illustrating a transcription dependence of the methylation patterning directed by MORC2 (**Figure 5F**). We also observed hypermethylation over zinc finger gene clusters that were specific to the MORC2 mutant but not CRISPRi organoids – in particular the ZNF genes hyperrepressed in pluripotent cells **(Figure 5G,H**). As major alterations of the methylation are present across the ZNF gene clusters (**Supp Fig 6C**), including gene bodies, we speculate that this could be driven by effects on 3D chromatin structure in MORC2 mutant cells, though this remains to be tested systematically. Together our observations demonstrate the critical role of MORC2 in directing transcription dependent CpG methylation patterning over its repetitive targets in early development.

**Figure 5.**
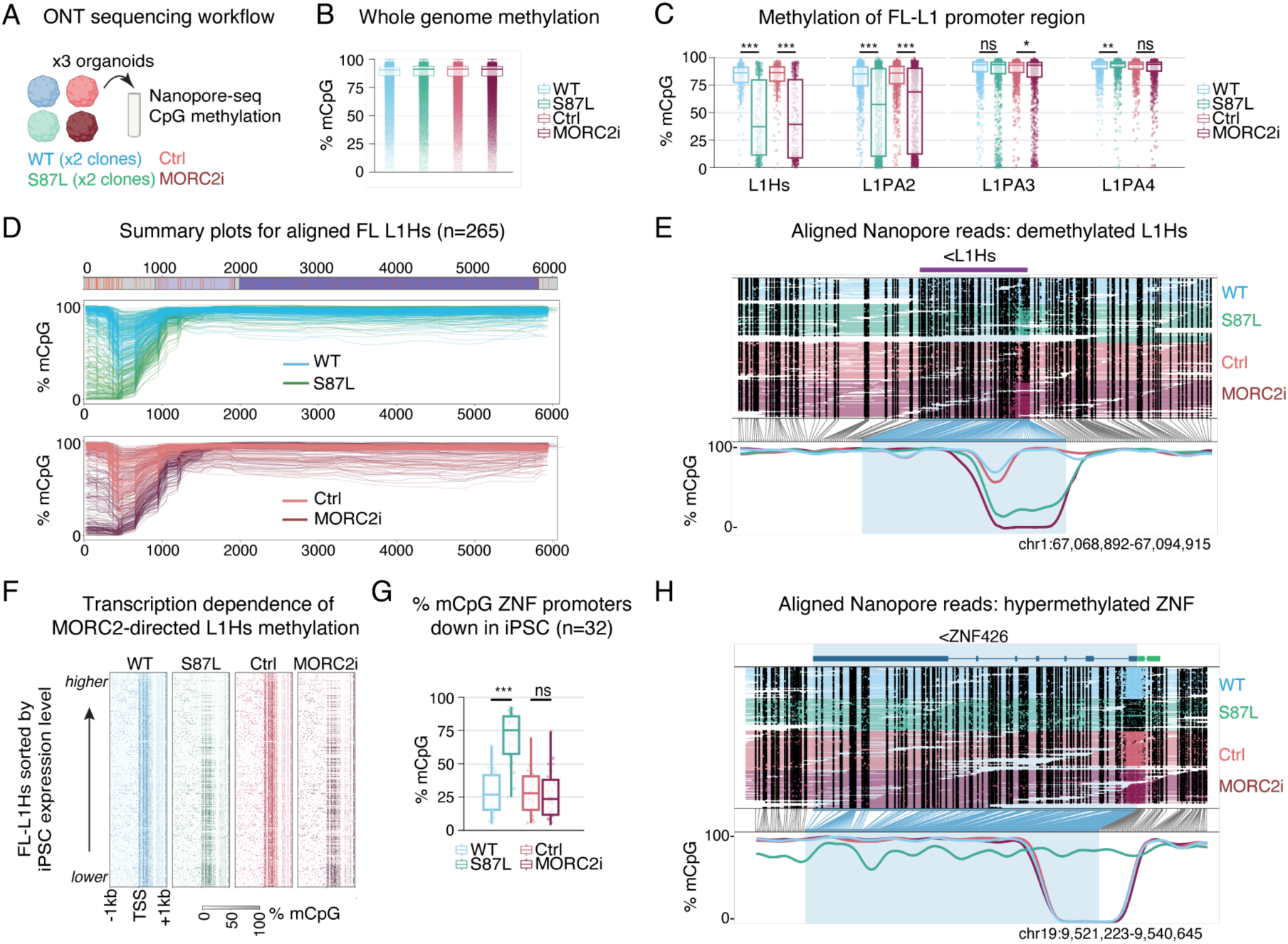
MORC2 directs CpG methylation patterning over the repetitive human genome. **(A)** Schematic of DNA methylation analysis workflow. For each clone and cell line, DNA from three organoids was extracted and pooled for Oxford Nanopore sequencing (ONT). **(B)** Boxplots showing the CpG methylation levels across the whole genome and **(C)** over the first 900-bp of full length (FL) evolutionary young LINE1 elements (L1HS, L1PA2-4), which corresponds to the 5’ UTR promoter region. P-value as calculated by a two-sided Wilcoxon rank sum test with continuity correction (p-values adjusted following Benjamini−Hochberg correction). **(D)** Composite plot showing CpG methylation levels of all reference FL-L1HS elements greater than 6 kb (hg38) over the entire element in the given conditions. **(E)** Aligned Nanopore reads showing loss of CpG methylation over the promoter of an example L1HS locus in S87L and MORC2i organoids. Methylated and non-methylated CpGs are represented by filled black circles and open (coloured) circles, respectively. **(F)** Heatmap showing the CpG methylation level over the promoter of FL-L1HS, sorted by expression level in hiPSC S87L mutants as assessed by promoter H3K4me3 level. Methylation changes are anti-correlated to their expression in hiPSCs. **(G)** CpG methylation level of the promoter of ZNF genes downregulated in MORC2-mutant iPSCs. **(H)** Aligned Nanopore reads showing hypermethylation over the promoter and general methylation alteration over the body of a hyperrepressed ZNF gene. ns, non-significant; *, p < 0.05; **, p < 0.01; ***, p < 0.001.

### L1 derepression upon exit of pluripotency causes loss of NPC proliferation

Finally we tested the impact of young L1 derepression on NPC differentiation. We returned to a directed 2D NPC protocol where both MORC2-CRISPRi and S87L mutant cells – i.e. those lines with high L1 expression – had undergone cell death in differentiation trials. Leveraging our recently-developed L1 CRISPRi lentiviral vector (**Fig. 6A,B**), which efficiently and specifically silences expression from full-length L1Hs, L1PA2 and L1PA3 subfamilies,^28^ we asked whether re-silencing L1s on the MORC2 S87L genetic background could rescue this apparent loss of cellular fitness. We generated six iPSC lines where WT or S87L clones were transduced with the L1 CRISPRi vector or a non-targeting control (**Fig. 6B**). We confirmed high young L1 expression in S87L clones by Western blot analysis of L1-orf1p, which was robustly lost upon L1 CRISPRi (**Fig. 6C**). Upon dual-SMAD inhibition to direct neural induction, we found that all lines exited pluripotency as assessed by RT-qPCR analysis of marker genes (**Supp Fig 8A**) and characteristic morphological changes of cellular polarization between days 4-7 of differentiation (**Fig. 6D, Supp Fig 8B**).^50^ By day 14, surviving cells from all lines expressed PAX6 and FOXG1 according to RT-qPCR and immunocytochemistry analysis (**Supp Fig 8A,B**). However, between day 9-14 of differentiation we observed striking cell death and 6-16 fold lower final cell yields in S87L clones transduced with the control CRISPRi vector, but not in those same clones transduced with the L1 CRISPRi vector, an effect that was reproducible across multiple blinded differentiation batches (**Fig 6E,F**). Thus, excessive L1 expression may be incompatible with rapid proliferation and/or early maturation of NPCs leading to loss of fitness, at least in this protocol. Taken together, our data indicate that MORC2-directed CpG methylation safeguards the somatic human genome against deleterious L1 expression upon exit of pluripotency and early neurodevelopment.

**Figure 6.**
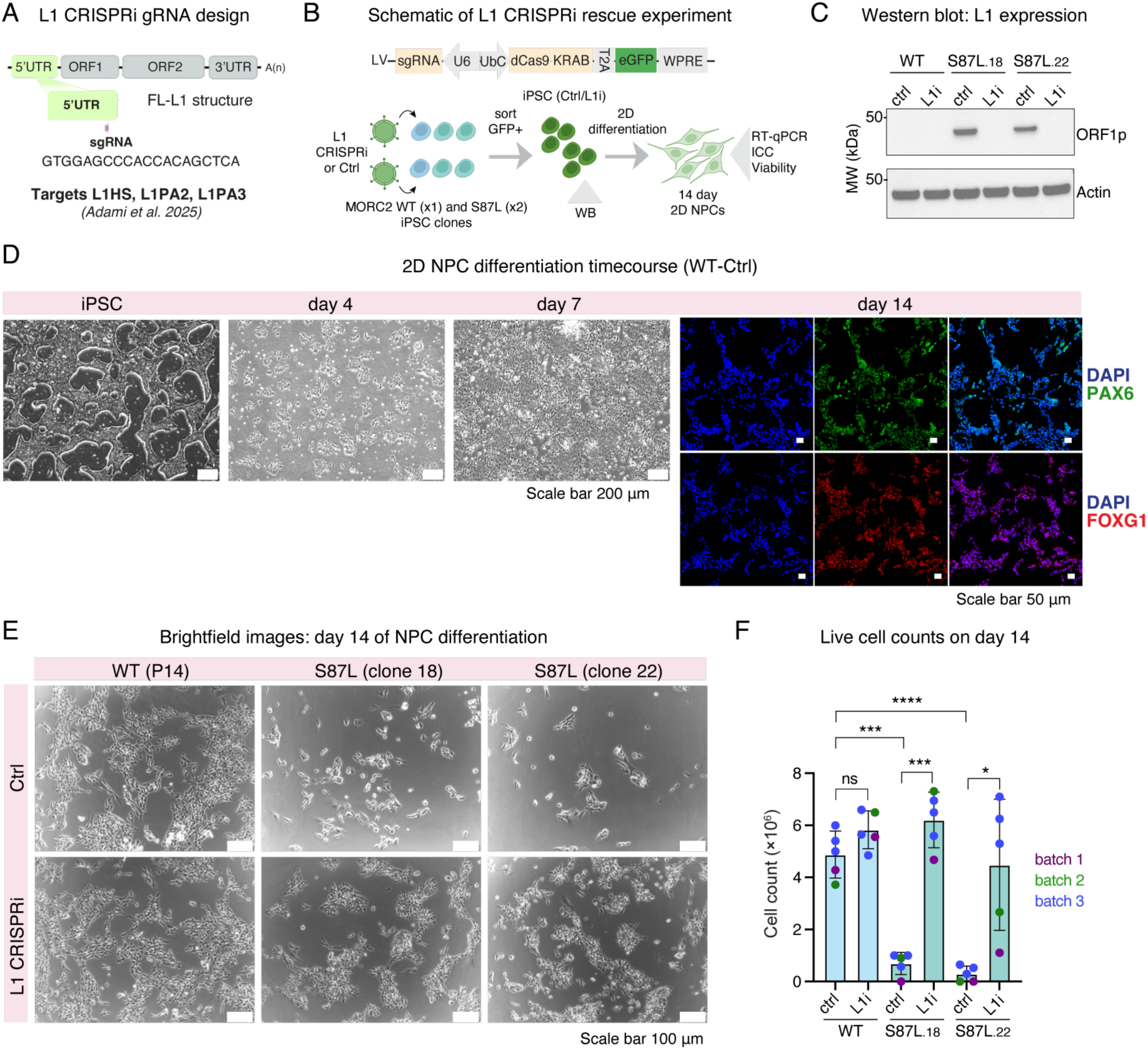
L1 derepression upon exit of pluripotency causes loss of NPC proliferation. **(A)** Schematic of L1 CRISPRi gRNA design to ectopically silence young full-length L1s. For full validation see Adami et al.^28^ **(B)** Experimental outline for L1 CRISPRi experiment in WT and S87L MORC2 iPSC lines, and subsequent 2D NPC differentiation. **(C)** Validation of young L1 silencing by Western blot of orf1p in the indicated iPSC lines. Experiment was repeated twice with similar results. **(D)** Representative brightfield (left) and ICC (right) images of WT Ctrl line during 2D NPC differentiation at the indicated timepoints. See also Supp Fig 8. **(E)** Representative brightfield images of the six indicated lines at the end of the 14-day 2D NPC differentiation protocol. **(F)** Quantification of live cell counts at the end of NPC differentiations from five replicates across three experimental batches (cells seeded on different days). Batch 3 was done in triplicate (cells seeded independently in different wells) to account for in-batch variation. Cell counts given are the mean value of two counts from each tube of cells. Bars indicate the mean (n=5) and error bars indicate the standard deviation. In batch 2, a small subset of cells were removed from all conditions at day 11 for subsequent immunocytochemistry. Experimenters were blinded to the identity of the cell lines during differentiation experiments. Significance was assessed using paired t-tests between indicated conditions. *, p < 0.05; ***, p < 0.001; ****, p < 0.0001; ns, not significant.

## DISCUSSION

In this study we report and relate two key mechanistic insights on the role of the MORC2 ATPase in epigenetic reprogramming of the repetitive genome. First, we find that reversible ATP-dependent dimerization is required for broad MORC2 accumulation over L1 retrotransposons, but not required for its binding to gene promoters, and that mutation of the ATPase module titrates MORC2 occupancy between these target classes. Second, we find that MORC2 chromatin binding directs CpG methylation patterns over its targets upon exit of pluripotency and early human neurodevelopment, which protects against the otherwise deleterious effects of unrestricted L1 expression. The model is summarized in **Figure 7**.

**Figure 7.**
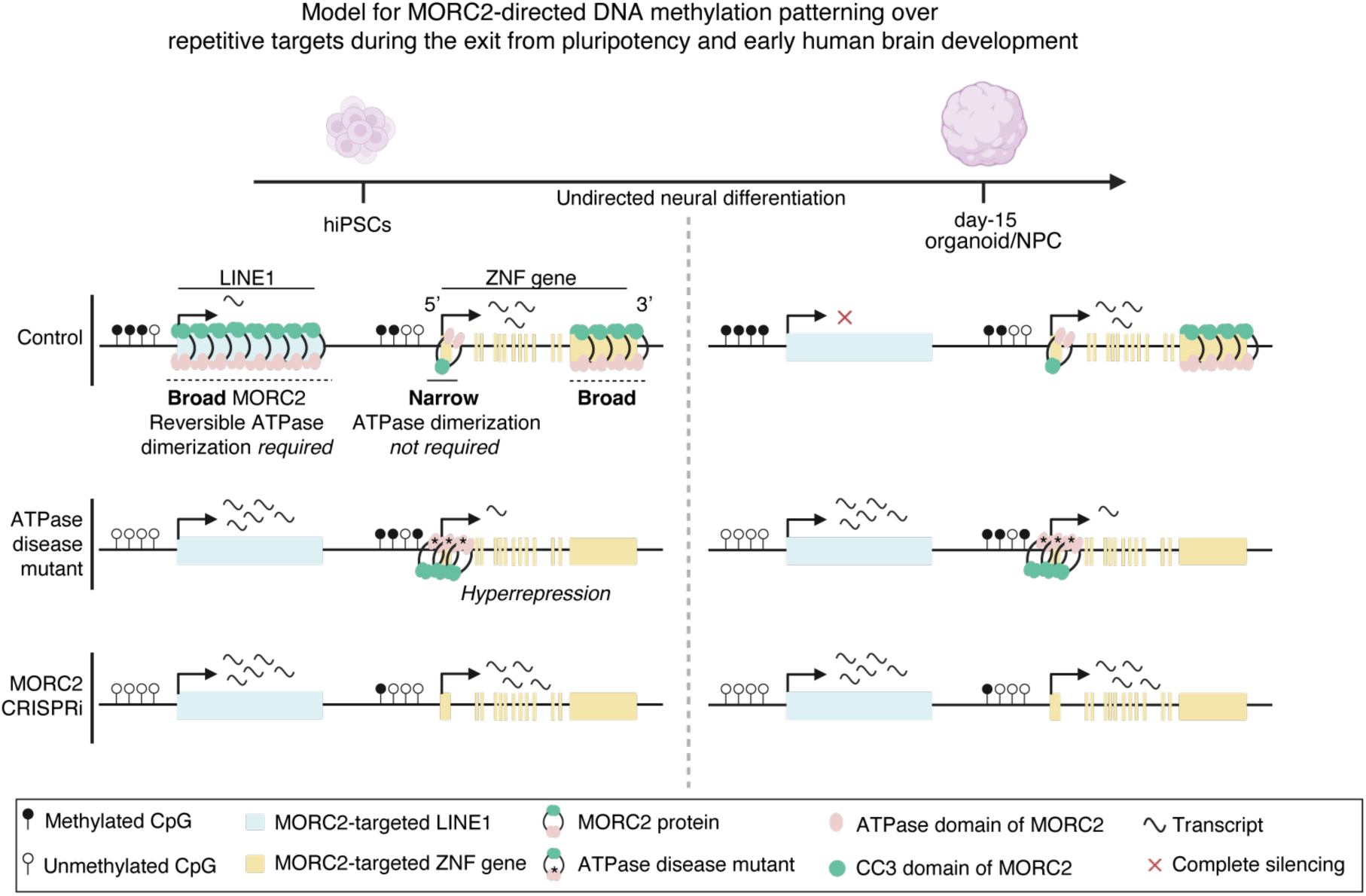
Model for MORC2-directed CpG methylation patterning over repetitive targets during the exit from pluripotency and early human brain development.

The physiological relevance of MORC2 ATP-dependent dimerization is evident from clinical genetics reports over the past ten years, where missense mutations in patients with severe neurodevelopmental disorders all map to the ATPase module.^45–47^ Our previous biochemical and structural studies identified that these mutations change the ATPase activity and dimerization dynamics of the purified protein,^37^ and here we provide a framework to explain how this translates at the level of chromatin binding. MORC2 has two main chromatin binding modes – broad accumulation over H3K9me3-marked sites, and narrow binding to gene regulatory elements marked by H3K4me3/H3K27Ac.^32^ When reversible ATP-dependent dimerization is perturbed by mutation of the ATPase module, MORC2 fails to accumulate over broad sites but binding is maintained or enhanced at narrow sites. Crucially, mutations that shift the equilibrium of N-terminal dimerization in either direction cause loss of function over L1s – supporting the notion that the lifetime of ATP-dependent MORC dimers is a critical parameter in driving chromatinization of certain targets.^36,37,39^ The observation that binding of MORC2 mutants to gene promoters is maintained or enhanced explains the hyper-repression phenotype observed in engineered cell lines or patient samples,^32,47^ and in the stem cell models of early brain development that we characterize here. That many of the genes hyperrepressed in MORC2 mutant cells encode KRAB-ZNFs, themselves associated with epigenetic TE silencing, highlights the central and multifaceted role of MORC2 in TE biology. Many of the hyperrepressed ZNFs in MORC2 mutant lines are transcription factors or repressors that each target hundreds of genomic locations, illustrating enormous potential for pleiotropic, indirect effects on chromatin and transcription of disrupting the balance of MORC2 chromatin binding sites via ATPase mutation. These complex gene regulatory networks require further investigation, but our data suggest that TE transcription and TE-induced chromatin structure are likely important molecular phenotypes to consider in MORC2 disorders.

Work by us and others has established that the HUSH-MORC2 corepressor targets transcribed and relatively hypomethylated repeats with little apparent preference for particular DNA sequence motifs.^8,33,51,52^ The prevailing model is that the complex is recruited to long, intronless units with some A-rich bias in the sense strand.^51^ In human pluripotent stem cells, the primary targets of MORC2 are the hominoid-specific LINE-1 retrotransposons that we recently reported to be transcribed and relatively hypomethylated at steady-state.^28^ This is likely the critical way that the HUSH-MORC2 axis preserves genome stability in early development: a subset of the youngest L1s remain competent for autonomous transposition and their encoded proteins provide the molecular apparatus for mobilization of other TEs. Transcription-dependent TE targeting is in contrast to the KRAB-ZNF/TRIM28 pathway, which is recruited to particular DNA sequence motifs in repeats via rapidly-evolving ZNF domains.^22–26^ Such complementary modes of targeting help explain why HUSH-MORC2 and KRAB-ZNFs may regulate overlapping yet distinct sets of genomic repeats and genes in different cellular contexts.^30,35^ The situation at the youngest human L1s is of particular interest: while numerous KRAB-ZNFs (and TRIM28) are competent to bind L1HS and L1PA2 subfamilies through conserved sequences in the L1 ORFs,^24,25^ deletion of 129 bp in the ancestral L1PA3 promoter region within the ZNF93 binding site may have evaded promoter methylation by this factor in evolutionarily-younger copies.^26,27^ Our data supports the idea that the HUSH-MORC2 corepressor is recruited to these youngest, transcriptionally-active elements for which a new KRAB-ZNF has yet to evolve.^51^

The interplay of these epigenetic silencing systems is of particular interest in early development, when methylation patterns of the germline are globally erased and re-established.^12^ The KRAB-ZNF/TRIM28 system is thought to protect against erasure of germline methylation at certain targets: for example, ZFP57 and ZFP445 bind DNA sequences at imprinting control regions, recruiting TRIM28 and DNMTs to maintain methylation at paternal or maternal imprints.^53,54^ Many TEs are also resistant to demethylation, but it is thought the youngest human L1s are demethylated early post-fertilization, maintaining relative hypomethylation throughout pre-implantation development.^27^ Although our primed iPSC models reflect a post-implantation developmental stage, we note that *MORC2* is also robustly expressed in the pre-implantation human embryo^55^ and our data suggest that the protein is a key factor that directs methylation over these youngest L1 copies. Whether MORC2 protects against erasure of germline methylation or promotes re-establishment of human L1 methylation following erasure is challenging to answer definitively, but the transcription-dependence of HUSH-MORC2 recruitment to L1s strongly favours a reestablishment model. Nevertheless, the underlying mechanism and timing of human L1 promoter methylation warrants further investigation, for example leveraging new blastoid models.^56^

Taken together, our study reveals how ATP-dependent dimerization of MORC2 is coupled to chromatin binding and DNA methylation patterning. The precise sequence of how establishment and/or maintenance DNA methyltransferases (DNMTs) are recruited to MORC2 targets in early development remains an interesting open question. MORC2-driven chromatin compaction is known to correlate with H3K9me3 deposition at certain sites,^8,32,33^ which may directly recruit maintenance methyltransferase DNMT1.^57,58^ However, since aberrant MORC2 mutant binding to certain (H3K9me3-negative) gene promoters also results in promoter hypermethylation, we suggest that MORC2 may be directly involved in licensing DNA methylation its target sites. This is somewhat reminiscent of a recent study in mouse gonocytes where Morc1, a germline-restricted TE repressor,^59^ was suggested to make distinct complexes leading to piRNA-directed H3K9me3 and DNA methylation at different TEs.^60^ Our data reinforce the model that the transcriptional activity of a locus determines its sensitivity to MORC2 binding and, we show here, targeted CpG methylation upon differentiation. Such a model could explain how the transcription factor yin-yang 1 (YY1) functions as both activator^61^ and repressor^31^ of human L1s: YY1 drive transcription initiation of these young elements in the pluripotent state, which marks the locus for MORC2-directed methylation. It is also noteworthy that certain young L1s are demethylated and transcriptionally-active in both organoids and human brain tissue.^3,28,31^ Further work is required to determine how MORC2-directed methylation may be avoided over these elements – or whether the elements are subject to active demethylation upon neural differentiation. How the mechanisms we describe here play out in terminally differentiated cells such as neurons also promises to be a fruitful area of research.

## METHODS

### Cell culture

#### hiPSCs

Two human iPSC lines were used in this study: Ctrl-14 hiPSCs, which were reprogrammed by the Cell and Gene Therapy Core at Lund Stem Cell Center from human fibroblasts via mRNA mediated reprogramming as described elsewhere,^62^ and a commercially-available line from RIKEN (RBRC-HPS0328 606A1). hiPSCs were cultured based on previously developed protocols.^50,63^ Briefly, cells were maintained in multi-well plates coated with Biolaminin 521 (0.7 mg cm^−2^), and fed daily with StemMACS iPS-Brew XF containing 0.5% penicillin–streptomycin (Gibco). hiPSCs were passaged upon reaching 70-80% confluency. Firstly, cells were rinsed with DPBS (Gibco), dissociated using Accutase (Gibco) for 5-7 min at 37°C then washed off by DMEM-F12 (Gibco) and KnockOut Serum Replacement (KSR) (Gibco) and centrifuged at 400g for 5 min. Rock inhibitor (10 μM, Miltenyi) was added to the media following cell thawing, lentiviral transduction, FACS and CUT&RUN experiments.

#### SH-SY5Y cells

SH-SY5Y human neuroblastoma cells (ATCC CRL-2266) were a gift from Harvey McMahon (MRC-LMB). Cells were cultured in flasks and fed every 3-4 days with complete culture medium containing DMEM-F12, 10% FBS, MEM non-essential amino acids solution and 1x penicillin/streptomycin (Gibco). When 80% confluent, cells were rinsed with DPBS (Gibco), incubated with trypsin (Gibco) for 5-7 minutes at 37°C to detach, and washed off with the media to quench the enzyme. To maintain a Mycoplasma-free culture, cells were routinely tested and returned negative.

#### Unguided cerebral organoid differentiation

Human cerebral brain organoids were generated using the STEMdiff Cerebral organoid kit (Stem Cell Technologies) following the manufacturer’s instructions and based on a previously established protocol.^64^ First, embryoid bodies (EB) were generated following disassociation and resuspension of hiPSCs in EB seeding medium (STEMdiff^TM^ Cerebral Organoids Basal Medium 1 and STEMdiff^TM^ Cerebral Organoids Supplement A, supplemented with 10 μM of Y-27632) in round bottom ultra-low attachment 96-well plates. On day 5, to avoid EB fusion individual EBs were distributed into single wells of ultra-low attachment 24-well plates and incubated with Induction Medium (STEMdiff^TM^ Cerebral Organoids Basal Medium 1 and STEMdiff^TM^ Cerebral Organoids Supplement B) for 48 h at 37°C. On day 7, organoids were then embedded into Matrigel (Stem Sell Technologies) and cultured in ultra-low attachment 6-well plates (10-12 organoids/well) with Expansion medium (STEMdiff^TM^ Cerebral Organoids Basal Medium 2, STEMdiff^TM^ Cerebral Organoids Supplement C, STEMdiff^TM^ Cerebral Organoids Supplement D). Up until day 15, organoids were fed twice (on day 10 &13) with Maturation media (STEMdiff^TM^ Cerebral Organoids Basal Medium 2, STEMdiff^TM^ Cerebral Organoids Supplement E). Organoids were harvested on day 15 for downstream analysis such as bulk RNA-seq (3 snap-frozen organoids per replicate), snRNA-seq (4-5 organoids per replicate), whole genome Oxford Nanopore sequencing (3-4 organoids per replicate), immunostaining (3-4 organoids) and CUT&RUN (3-4 organoids per replicate).

#### 2D NPC differentiation

Transduced iPSC cell lines vectors were differentiated to neural progenitor cells (NPCs) using a 2D-protocol based on dual inhibition of the SMAD-signalling with Noggin and SB431542 from the start of the differentiation as described elsewhere.^50^ A two-day coating protocol was followed. On the first day, 6-well plate was coated with 100 μL of poly-L-ornithine (PLO, Sigma-Aldrich) diluted in 10 mL of PBS +/+ (1.2 mL per well) and incubated at 37℃ overnight. The following day, PLO solution was aspirated and the wells were washed three times with PBS +/+. PBS +/+ was aspirated and the plate was coated with 50 μL of Laminin-23017-015 (Fisher Scientific) diluted in 10 mL of PBS +/+ (1.2 mL per well). The plate was wrapped in parafilm and placed in the fridge to incubate at 4℃ overnight. Before seeding the cells, Laminin-23017-015 was aspirated, and the wells were washed once with PBS +/+. hiPSC lines were counted and 90,000 cells were transferred to a tube, pelleted then resuspended in N2 medium supplemented with SB431542, Noggin and Rock Inhibitor, before being seeded into the wells. On days 2, 4 and 7 of the differentiation, N2 media supplemented with SB431542 and Noggin was supplied to the cells. On the 9th and 10th day of differentiation N2 media without supplements was used. On day 11 cells were passaged and replated in B27 media (Table 3) supplemented with brain-derived neurotrophic factor (BDNF, R&D systems), ascorbic acid (AA, Sigma-Aldrich) and Revita (Thermo Fisher Scientific). B27 media supplemented with BDNF and AA was supplied to the cells every day until the day of the harvest.

**Table 1.**
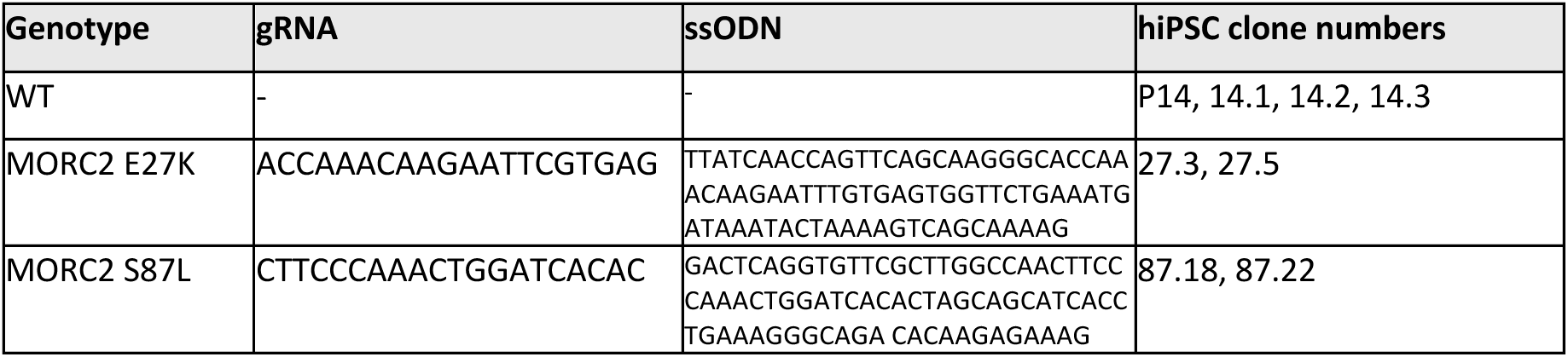
CRISPR editing of *MORC2*.

**Table 2.**
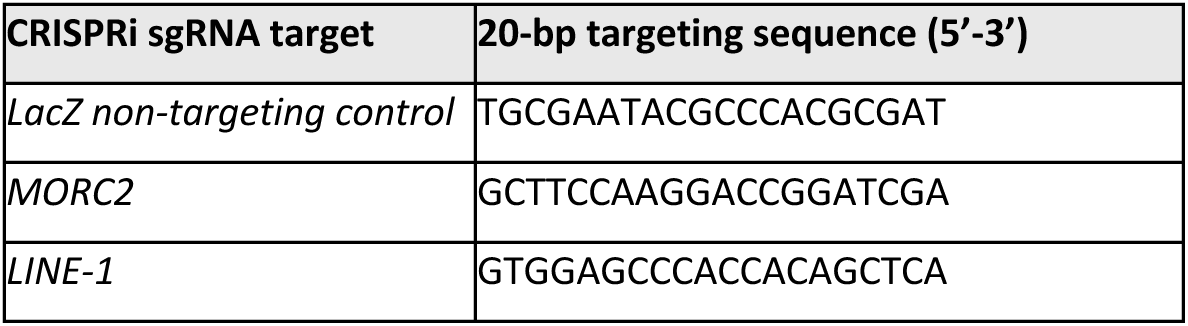
CRISPRi gRNA sequences.

**Table 3.**
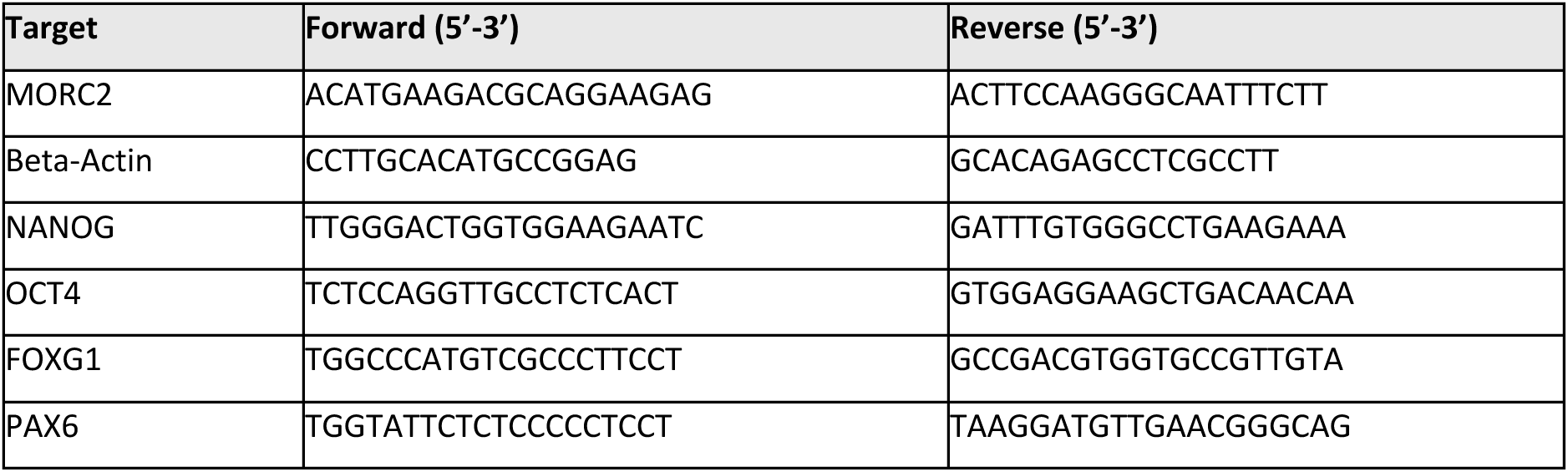
RT-qPCR primer sequences.

### Lentiviral vectors

Lentiviral vectors were produced according to a standard protocol described previously.^65^ Briefly, HEK293T cells with a confluency of ∼80% were co-transfected with three packaging plasmids (pMDL, psRev, and pMD2G) and the relevant transfer plasmid using polyethyleneimine (PEI; Polysciences), in DPBS. Cell culture media containing lentiviral particles was harvested 48 h after transfection, filtered, and centrifuged at 25,000xg for 1.5h at 4 °C. After removing the supernatant, pellets were gently resuspended in cold PBS and incubated overnight at 4°C. Finally, the lentivirus was aliquoted and stored at -80 °C until use. Lentiviral titration was determined using qPCR targeting the viral WPRE sequence and comparison to a reference virus batch with known titre.

### CRISPR approaches

#### CRISPR KO

To generate MORC2 KO SHSY5Y cells two sgRNA were cloned into pSpCas9(BB)-2A-Puro vector (Addgene #48139). sg21 (GGGATCAATATAGAGCACAG) targets exon 9 and sg22 (GCGGACATTAGAAGTACGCCT) exon 11 of *MORC2*. 8 mg of the plasmids was electroporated into 2x10^6^ cells using 100ul tips from Neon electroporation system and placed into 25 cm flasks. After 24 h, 4 mg/mL of puromycin was added to select for electroporated cells. After 10 days single cells were sorted into 96-well plates. Chosen clones were first analysed for the presence of the MORC2 band using Western blotting. Cells were lysed for 30 min at 4°C in buffer containing 300 mM NaCl, 100 mM Tris pH 8.0, 0.2 mM EDTA, 0.1% NP40, 10% glycerol supplemented with protease inhibitors. Lysates were cleared by centrifugation and supernatant was used for Western blotting using MORC2 (anti-rabbit MORC2, Bethyl), and actin antibodies (Mouse α-β-actin, ab8226). DyLight-680 or 800-conjugated secondary antibodies (Thermo Fisher Scientific) were used at 1:10,000 dilution and blots were imaged on the Odyssey near-infrared system (LI-COR). Following isolation of genomic DNA (Qiagen), the knockout was confirmed using Sanger sequencing and TIDE analysis.

#### CRISPR-editing

To introduce MORC2 S87L and E27K mutations in Ctrl-14 hiPSCs, HiFi Cas9v3, relevant gRNAs and desired sequences to target the MORC2 gene were delivered to cells as Ribonucleoprotein (RNP) and single-stranded oligodeoxynucleotide (ssODN) via nucleofection (P3 Primary Cell 4D-Nucleofector X Kit, Lonza) (**Table 1**). CRISPR editing efficiency was estimated in bulk using genomic PCR and restriction enzyme digestion followed by clonal expansion. Individual clones were collected and primarily tested for the presence of the edit using genomic PCR and restriction enzyme digestion followed by Sanger sequencing. The selected clones with the desired edits underwent karyotyping and sequencing of the 5 top-predicted off-targets. Once these quality controls were passed, immunostaining of pluripotency markers OCT4, NANOG and SOX2 was performed on selected clones.

#### CRISPR interference

A previously-described protocol for CRISPR interference was used to silence *MORC2* or young L1 transcription in hiPSCs.^66^ gRNAs (**Table 2**) were cloned into a dCas9-KRAB-T2A-GFP lentiviral backbone containing both the gRNA under the U6 promoter and dCas9-KRAB-T2A-GFP under the Ubiquitin C promoter (pLV hU6-sgRNA hUbC-dCas9-KRAB-T2a-GFP) using annealed oligos and the BsmBI cloning site. An identical CRISPRi vector containing a gRNA sequence absent from the human genome (LacZ) was used as a non-targeting control. hiPSCs were transduced with MOI 5 and GFP-positive cells were sorted using FACS (FACSAria, BD sciences) on day 10 at 10°C. Sorted cells were centrifuged at 400x*g* for 7 min and pellets were snap frozen on dry ice and kept at −80°C for downstream analysis, or maintained in culture. MORC2 CRISPRi was validated in Pandiloski et al.^33^. Young LINE-1 silencing was validated in Adami et al.^28^

### RT-qPCR

RT-qPCR was used to check the efficiency of MORC2 CRISPRi. 500 ng RNA was extracted from hiPSCs and used for reverse transcription to generate cDNA with random hexamer primers and the Superscript III reverse transcriptase (Invitrogen). RT-qPCR was performed using SYBR Green I master (Roche) on a LightCycler 480 instrument (Roche). Data were analysed with the ΔΔCt method normalized to housekeeping gene *ACTB*. Primer sequences are given below.

### Lentiviral expression of HA-MORC2 variants

To express WT and mutant versions of MORC2 on the SHSY5Y MORC2-KO background, we cloned MORC2 into the lentiviral pHRSIN-pSFFV-WPRE-pPGK-Hygro vector (a gift from Paul Lehner, University of Cambridge) with an N-terminal HA-tag. For lentiviral production 293T cells were co-transfected with lentiviral MORC2 vector, pCMVΔR8.91 and pMD.G using the TransIT-293 transfection reagent (Mirus). Supernatants were collected two days post-transfection and cleared by filtering through 0.45 mm membranes. 1x10^6^ of cells were transduced by centrifugation for 90 min at 1000xg in the presence of 300 μL lentivirus-containing supernatant and 5 μg/mL of polybrene. The following day, cells were transferred into 10cm dishes and hygromycin at 400 μg/mL added for selection.

### Quantitative genotyping qPCR

Quantitative genotyping PCR (qgPCR) was performed to rule out unintended CRISPR-induced on-target effects (onTEs) at the edited sites. We followed the protocol described in Weisheit et al., 2021.^67^ In brief, genomic DNA was extracted from iPSC cells using the DNeasy Blood & Tissue kit (QIAGEN). IDTPrimerQuest was used to design primer pairs to amplify the regions surrounding the edits and FAM-labeled target-specific probes and purchased from IDT (see **Table 4**). We use the TERT Taqman Copy Number Reference assay (Thermo Fisher Scientific) labeled with VIC as a control assay. The qgPCR reaction mix was prepared using the Prime-Time Gene Expression Master Mix (IDT). Each sample was run in technical triplicates on a LightCycler 480 instrument (Roche). Protocol guidelines were followed to perform the Ct values analysis and total allele number calculation.

**Table 4.**
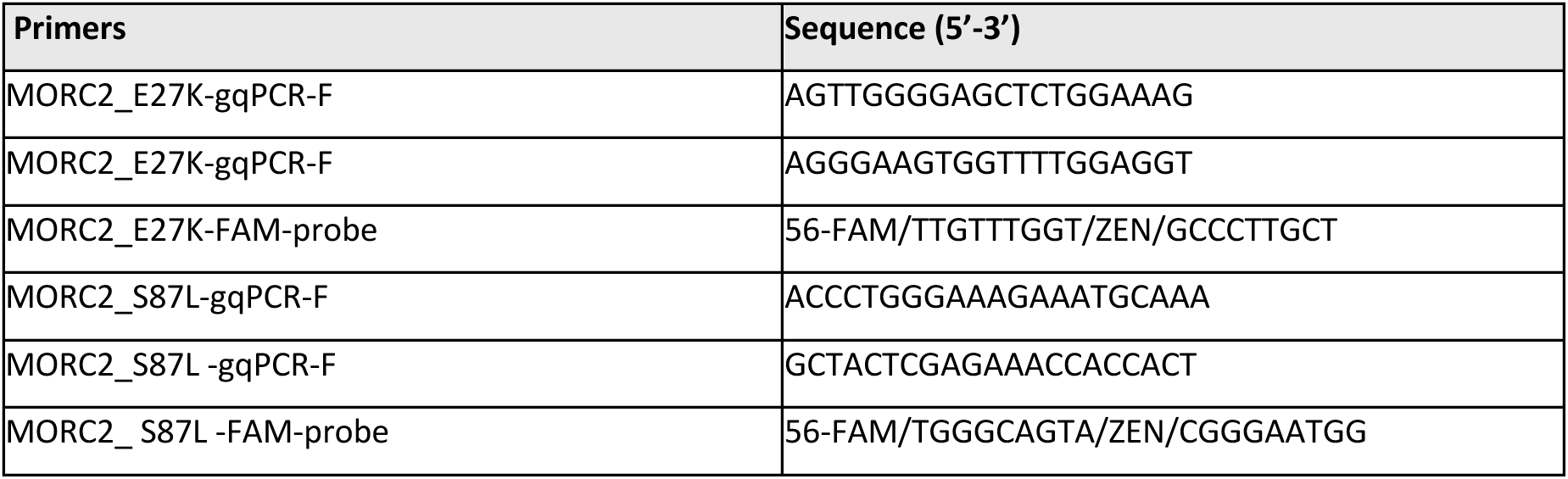
Primers for quantitative genotyping qPCR.

### Bulk RNA sequencing and analysis

RNA was extracted from cells and organoids using the RNeasy mini and QIAshredder kits (Qiagen) respectively. Sequencing libraries were prepared using the Illumina TruSeq Stranded mRNA Library prep kit optimized for long fragments and subsequently sequenced on a NovoSeq X Plus platform with paired end reads (2x150bp) sequencing.

#### Genes and repetitive element subfamilies quantification

The generated reads were mapped to the human reference genome hg38 (GRCh38) with STAR aligner (v 2.6.0)^68^ with modified parameters (--outFilterMultimapNmax 100 and --winAnchorMultimapNmax 200). BAM files were sorted using SAMtools (v 1.16.1).^69^ For visualisation, BigWig files were created from the BAM files using bamCoverage (v 2.5.4) applying RPKM normalization. Gene quantification was conducted with Subread featureCounts (v1.6.3; flags -p and -s 2 for paired-end reads and strand-specific reads, respectively) using the reference annotation GENCODE version 38. Repetitive elements quantification at a subfamily level was conducted with TEtranscripts (v 2.2.3) using default options and using a curated annotation file.^70^

#### Individual repetitive elements quantification

To obtain an accurate quantification of individual repetitive elements, alignment of the reads to the reference genome with STAR aligner was conducted to allow only for uniquely mapped reads. To this end, the reads were allowed to map only to a single locus (--outFilterMultimapNmax 1) and the allowed mismatched ratio was decreased (--outFilterMismatchNoverLmax 0.03) compared to default settings. After sorting the obtained BAM files with SAMtools, the quantification was conducted with featureCounts (v1.6.3) using the curated annotation file provided within the TEtranscripts package. Generation of the BigWig files was conducted as described above.

#### Differential expression analysis

Differential expression analysis was conducted with DESeq2 (v 1.40.2)^71^ using as input the count matrices obtained from featureCounts and TEtranscripts. Raw reads were normalized with DESeq2’s median-of-ratios normalization method and fold changes were shrunk using DESeq2’s lfcShrink. For creating the heatmaps, the mean of replicates for each condition + 0.5 was used and further visualized with pheatmap.

### Nuclear isolation

Nuclei extraction was performed from day-15 organoids according to previously described protocols.^72,73^ Briefly, organoids were dissociated in cold lysis buffer (0.32 M sucrose, 5 mM CaCl2, 3 mM MgAc, 0.1 mM Na2EDTA, 10 mM Tris-HCl, pH 8.0, 1 mM DTT) using a 1ml tissue douncer (Wheaton). Isolation was sucrose gradient-based, thus the lysate was layered carefully on top of a sucrose cusion (1.8 M sucrose, 3 mM MgAc, 10 mM Tris-HCl pH 8.0, and 1 mM DTT diluted in milliQ water) and then centrifuged for 2h and 15 minutes at 30,000xg 4 °C. Supernatant was thoroughly aspirated, pellet was soften for 10 minutes by adding nuclear storage buffer (15% sucrose, 10 mM Tris-HCl pH 7.2, 70 mM KCl, and 2 mM MgCl2, all diluted in milliQ water), resuspended in dilution buffer (10 mM Tris-HCl pH 7.2, 70 mM KCl, and 2 mM MgCl2, diluted in milliQ water) and filtered using a 70um cell strainer. Finally, nuclei were sorted using FACS (Aria, BD Biosciences) at low flow rate and 4° C.

### Single-nuclei RNA-sequencing and analysis

Approximately 8,500 to 10,000 nuclei per sample were prepared for single-nucleus RNA sequencing and loaded onto the Chromium Next GEM Chip G using the Single Cell Kit, along with the reverse transcription mastermix, following the manufacturer’s protocol for the Chromium Next GEM Single Cell 3’ Kit (10X Genomics, PN-1000268). Single-cell gel beads in emulsion (GEMs) were generated during this process. cDNA was then amplified according to guidelines from 10X Genomics, using 13 PCR cycles for the 3’ libraries. Sequencing libraries were constructed with unique dual indices (TT set A), pooled, and sequenced on a NovaSeq X Plus system using a 100-cycle kit with 28-10-10-90 read lengths. Cell Ranger count (v 6.0.0) was run with default settings,^74^ with an mRNA and pre-mRNA reference for single-cell samples (generated using 10x Genomics Cell Ranger 6.0.0 guidelines) for single-nucleus samples. Samples clustering was analyzed using Seurat (version 5.1.0).^75^ Each sample was filtered according to two criteria, I) nuclei with percentage of mitochondrial content over 2% (perc_mitochondrial) and II) nuclei with number of features (n_Feature) lower than 600 and greater than 2 standard deviations were filtered out. The counts were normalized using Seurat’s LogNormalize (NormalizeData) and clusters were defined with a resolution of 0.1 (FindClusters). Standard Seurat functions (e.g. DimPlot and DotPlot) were used for all downstream visualization of the data.

### Oxford Nanopore sequencing and analysis

For each sample, high molecular weight DNA was extracted from pooled organoids (n = 3) using the Nanobind HMW DNA extraction kit (PacBio) according to manufactureŕs instructions. Extracted DNA was then depleted of short DNA fragments (< 10 kb) and sheared to 10 kb using the Short Reads Eliminator (SRE) XS kit (PacBio) and the g-TUBE (Covaris), respectively. The sequencing libraries for LacZ control and MORC2 CRISPRi samples were prepared using the Ligation Sequencing Kit V14 (SQK-LSK114 - ONT), while for the WT control (clones P14 and 14.3) and S87L MORC2 mutant (clones 87.18 and 87.22) the library was prepared, and the clones were barcoded and using the Native Barcoding Kit 24 V14 (SQK-NBD114.24 - ONT). All libraries were sequenced on FLO-PRO114M flow cells on a PromethION device (ONT) for 72 h. Basecalling and alignment to the human reference genome hg38 was conducted with Dorado (v0.7.1) using the DNA model dna_r10.4.1_e8.2_400bps_sup@v4.3.0 and the DNA modification model dna_r10.4.1_e8.2_400bps_sup@v4.3.0_5mCG_5hmCG@v1 for the detection of CpG methylation. Resulting BAM files were sorted with SAMtools (v1.18). Methylartist (v1.3.0) was used for the subsequent analysis of the methylation data visualised through Methylartist and R (v4.4.2).^76^ Modkit (v0.3.3) and BedGraphToBigWig (v4) were used to produce bedGraph from the BAM files and BigWigs from the bedGraph files, respectively. The BigWig files were then used for visualisation of the methylation with computeMatrix and plotHeatmap (DeepTools v2.5.4).

### CUT&RUN

Two different CUT&RUN approaches were followed. For profiling histone modifications (H3K4me3, H3K9me3, H3K27Ac), a standard CUT&RUN protocol was used. To profile MORC2 chromatin binding, a crosslinked protocol was used.

#### Standard CUT&RUN

The CUT&RUN protocol used here has previously been described.^77,78^ Briefly, 500k cells were harvested as described above (see cell culture: hiPSCs or SHSY5Y) and washed three times using Wash buffer containing 20 mM HEPES pH 7.5, 150 mM NaCl, 0.5 mM spermidine, 1x Roche cOmplete protease inhibitors. Cells were then incubated and attached to 10 uL pre-activated ConA-coated magnetic beads (Bangs Laboratories) on an end-to-end rotator for 7-10 min. To activate ConA-coated magnetic beads, they were washed twice with binding buffer containing 20 mM HEPES pH 7.9, 10 mM KCl, 1 mM CaCl2, 1 mM MnCl2. Bead-bound cells then were incubated with primary antibody solution (20 mM HEPES pH 7.5, 0.15 M NaCl, 0.5 mM Spermidine, 1x Roche complete protease inhibitors, 0.05% w/v digitonin, 2 mM EDTA and 1:100 primary antibody) overnight at 4 °C. On the second day, bead-bound cells were washed twice using digitonin buffer (20 mM HEPES pH 7.5, 150 mM NaCl, 0.5 mM Spermidine, 1x Roche cOmplete protease inhibitors, 0.05% digitonin) then incubated with pAG-MNase enzyme resuspended in digitonin buffer at 4 °C for 1 h with a gentle mixing. Bead-bound cells were then washed twice with digitonin buffer, resuspended in 100μL digitonin buffer, chilled to 0 °C for 5 min then incubated for 30 min at 0 °C with 2 mM CaCl2 to activate genome cleavage. Finally, the reaction was quenched by adding 100 mL 2x stop buffer (0.35 M NaCl, 20 mM EDTA, 4mM EGTA, 0.05% digitonin, 50 ng/mL glycogen, 50 ng/mL RNase A) and gentle vortexing followed by a 30 min incubation at 37 °C to release the genomic fragments. The supernatant was collected and purified using PCR clean-up spin columns (Macherey-Nagel).

#### Day-15 organoid CUT&RUN

3-6 organoids per cell line was used. Organoids were collected on day-15 of differentiation, washed with PBS, and incubated 3 minutes with Accutase on a shaker at 37 °C to dissociate cells. They were then gently resuspended in Accutase and incubated on the shaker for another 4 minutes followed by a 5-minute centrifugation at 400xg. Finally, the cell suspension was filtered to remove Matrigel. The standard *CUT&RUN* protocol was followed as described above.

#### Crosslinked CUT&RUN

The approach was same as above with addition of the followings: a cross-linking step, an increase in cell number and a slightly higher centrifugation force. 900K-1M cells were collected by centrifugation at 600xg for 3 minutes and crosslinked for 1 min using 1% formaldehyde (in cell culture media, diluted from a 16% solution, Thermo Scientific). Cross-linked cells were then washed three times with wash buffer containing 20 mM HEPES pH 7.5, 150 mM NaCl, 0.5 mM spermidine, 1x Roche complete protease inhibitors, 1% Triton X-100, 0.05% SDS. To avoid loss of the crosslinked cell/nuclear pellet during washes, a slight increase in centrifugation speed (1000-1300xg) was applied. Cells were incubated with ConA beads for 7-10 minutes as above and then resuspended in primary antibody buffer (20 mM HEPES pH 7.5, 150 mM NaCl, 0.5 mM spermidine, 1x Roche complete protease inhibitors, 1% Triton X-100, 0.05% SDS, 0.05% digitonin, 2 mM EDTA). After an overnight incubation at 4 °C with the primary antibody, bead-bound cells were thoroughly washed two times with digitonin buffer (20 mM HEPES pH 7.5, 150 mM NaCl, 0.5 mM Spermidine, 1x Roche complete protease inhibitors, 1% Triton X-100, 0.05% SDS, 0.05% digitonin). Following the 37 °C incubation to release genomic fragments, 0.09% SDS and 0.22 mg/ml proteinase K (Thermo Scientific) was added to supernatant to reverse the crosslinking reaction. Finally, DNA was purified using PCR clean-up spin column (Macherey-Nagel).

### CUT&RUN sequencing and analysis

Hyperprep kit (KAPA) with unique dual indexed adaptors were used to prepare sequencing libraries which were then pooled and sequenced on NovaSeqX plus machine (Illumina). Paired-end reads (2×75) were aligned to the human genome (GRCh38) using bowtie2 (v 2.4.5) (–local –very-sensitive-local –no-mixed –no-unal –no-discordant –phred33 -I 10 -X 700),^79^ converted to BAM files and further sorted with SAMtools (v 1.16.1). Peaks were called using HOMER (v 4.10). Tag directories were created from the BAM files using makeTagDirectory, and peaks were called with findPeaks (-style histone -minDist 1000 and - size 300). When available, MORC2 CRISPR KO samples were used as control for peak calling within the findPeaks command (-i). For the NPC lines, where the MORC2 CRISPR KO was not available, the IgG sample was used as control, and MORC2 peaks with signal in the IgG samples were removed. Peaks were filtered based on number of tags (> 10) and on fold-change compared to control (> 8).Reads per kilobase per million mapped reads (RPKM)-normalized BigWig coverage tracks were made with bamCoverage (DeepTools, v 2.5.4). Heatmaps matrices were created by computeMatrix (DeepTools v2.5.4) using called peaks as the input regions and RPKM normalized bigwigs as input signal. Matrices were then visualized using plotHeatmap (DeepTools, v2.5.4), adjusting the upstream and downstream visualization parameters (-a and -b) and splitting the signal in heatmaps (--kmeans 2).

### Immunostaining

#### hiPSCs immunostaining

Cells cultured on 24-well ibidi plates were rinsed three times with DPBS and fixed in 4% formaldehyde for 15 min at room temperature. Before blocking with TKPBS (KPBS with 0.25% Triton X-100) and 5% Normal Donkey Serum (NDS), cells were washed another three times with DPBS. After 1h blocking, they were incubated overnight at 4 °C with primary antibody or TKPBS + 5% NDS for negative controls. The next day cells were rinsed in TKPBS twice for 5 minutes each and once in TKBPS/ 5% NDS. They were then incubated with secondary antibody solution (TKPBS + 5% NDS) at room temperature for 2 h followed by a 5-minute incubation with DAPI (1:1000, Sigma D817) diluted in TKPBS. After two washes with KPBS, cells were imaged using a Zeiss 780 confocal microscope.

#### Day-15 organoid immunostaining

Organoids harvested at day 15 of differentiation were gently rinsed once with DPBS (Gibco) in 24-well plates then fixed with 4% paraformaldehyde (PFA) for 90-120 minutes at room temperature. After fixation, organoids were washed three times with KPBS for 10 minutes each time and incubated overnight with 30% sucrose at 4 °C overnight on gentle shake. The following day, organoids were immersed in OCT (catalog no. 45830, HistoLab), snap frozen on dry ice and stored in -20°C until use. Before staining, organoids were sliced into 20-μm sections using a cryostat (Thermo Scientific Cryostar NX70) at -20°C and placed onto Superfrost Plus microscope slides. Borders of each slide were defined using a hydrophobic PAP Pen before starting the immunostaining. After two washes with KPBS, slides were incubated with blocking solution (TKPBS & 5% NDS) for 1 h at room temperature. They were then incubated with primary antibody solution (relevant primary antibody diluted in TKPBS & 5% NDS) overnight at 4 °C. The following day slides were washed three times with KPBS and incubated with secondary antibody solution for 1 h at room temperature. Finally, after three more washes, slides were incubated with DAPI, rinsed off with KPBS, mounted using FluorSave mounting medium and imaged with a Zeiss 780 Confocal microscope. Further image adjustments were performed in batch on Image J.

### STORM

Cells were seeded on glass 8-well Ibidi dishes and next day fixed with 4% formaldehyde in PBS for 15 min. Membranes were permeabilised with 0.1% Triton-X100 in PBS for 5 min. For staining, wells were first blocked with 10% FBS in PBS then incubated with primary anti-HA antibody at 1/500 followed by secondary Alexa Fluor 647 at 1/1000. After washing samples were fixed again for 5 min, washed and kept in PBS at 4°C till the imaging. STORM data was acquired using Nikon N-STORM inverted microscope with 647 nm laser line, Apochromat TIRF 100×/1.49 NA oil immersion objective and single-photon detection iXon Ultra DU897 EMCCD camera (Andor). Samples were washed with PBS and then STORM buffer (7mM NaOH, 100 mM mercaptoethylamine (MEA) in PBS) was added for the imaging. Samples were illuminated with the 647 nm laser at 2–3 kW·cm^−2^ and 100 000 frames were collected for each field of view at a frame rate of 50–70 Hz. Data was processed in NIS elements Advanced Research with N-STORM module. Further image processing and cropping was performed using ImageJ.

### Western blot

#### MORC2 KO SHSY5Y cells and complementation

Cells were lysed for 30 min at 4°C in buffer: 300 mM NaCl, 100 mM Tris pH 8.0, 0.2 mM EDTA, 0.1% NP40, 10% glycerol supplemented with protease inhibitors. Lysates were cleared by centrifugation and supernatant used for Western blotting, probed with indicated antibodies and scanned with a near-infrared system (LI-COR Odyssey) after incubation with DyLight 680-or 800-conjugated secondary antibodies (Thermo Fisher Scientific, 1:10,000 dilution).

#### Assessment of MORC2 and L1 expression in iPSCs

Cell lysis was performed using either RIPA (Sigma-Aldrich) containing complete protease inhibitor cocktail or 1:100 benzonase in 1% SDS on ice for 30 minutes and pelleted for 20 minutes at 4 °C and 12,000x*g*. Before running on a 4–12% Tris-glycine SDS-PAGE gel (200 V, 40-45 min, RT), the supernatant was aspirated into new tubes and mixed with loading dye (Novex LDS 4x) and reducing agent (Thermo) then boiled for 10 minutes at 70 °C. To transfer proteins from gel to membrane, premade transfer stacks (iBlot™ 2 NC Mini Stacks, Invitrogen™) were used in the iBlot™ 2 gel transfer device, using the P0 preset template, according to manufacturer’s instructions. The membrane was briefly rinsed with TBS-T (Tris-buffered saline with 0.1% Tween) and blocked in 5% skimmed milk diluted in TBS-T (MTBST) for 1h at RT on a shaker. Membrane was rinsed once in TBS-T and incubated at 4 °C overnight with primary antibody diluted in MTBST. The following day membrane was washed three times in TBS-T each time for 5-10 minutes with shaking. It was then incubated with HRP-conjugated anti-mouse (Cell signalling) or HRP-conjugated anti-rabbit secondary antibody (Santa Cruz Biotechnology, 1:5,000) diluted in MTBST. Before protein detection, membrane was washed three times in TBS-T each time for 5-10 minutes with shaking and once with TBS. Protein was detected by chemiluminescence using ECL Select reagents (Cytiva) according to manufacturer’s instructions and imaged on a Chemi-Doc system (BioRad). Finally, β-actin staining was performed by stripping the membrane using the Restore PLUS Western Blot Stripping Buffer (Thermo) as per instructions, re-blocking for 1 h in MTBST, followed by incubation with HRP-472 conjugated anti-β-actin (1:50000 dilution Sigma A3854). For uncropped blots, see **Supp Fig 9.**

### Antibodies

*CUT&RUN*: Goat anti-rabbit IgG (abcam ab97047), Rabbit anti-H3K4me3 (Active Motif 39159), Rabbit anti-MORC2(A300-149A, Bethyl), Rabbit anti HA-Tag (C29F4) mAb 3724, Rabbit anti-H3K9me3 (abcam 8898), Rabbit anti-H3K27ac (MABE647 clone RM172) (1:50-1:100 dilution). *Immunostaining*: Rabbit anti-NANOG (1:600 dilution, Abcam, RRID: AB446437), Mouse anti-OCT3/4 (1:300 dilution,Santa Cruz, RRID: AB628051), Rabbit anti-Sox2 1:500 (Merck AB5603, RRID: AB2286686), DAPI (1:1000 dilution, Sigma-Aldrich), Mouse anti-ORF1p, clone 4H1 (1:300 dilution, Sigma-Aldrich, RRID: RRID:AB2941775), Rabbit anti-PAX6 (1:300, Biolegend, 901301), Mouse anti-ZO1 (1:300, Invitrogen, 339100), Mouse anti-HA11 (Biolegend, 901501), Donkey anti-mouse Cy3 (1:500 dilution, Jackson Lab), Donkey anti-rabbit Alexa647 (1:500 dilution,Jackson Lab), Alexa-488 anti-mouse (Thermofisher, A21202, RRID AB141607), Alexa-568 anti-rabbit (Thermofisher A10042, RRID AB2534017).

*Western Blot*: Rabbit anti-MORC2 (A300-149A, Bethyl), Mouse anti-HA (HA.11 Biolegend 901501, clone 16B12), Anti LINE-1 ORF1p clone 4H1 (Merck Millipore, MABC1152), HRP-472 conjugated anti-β-actin (1:50000 dilution Sigma A3854), HRP-conjugated anti-mouse IgG (1:10000, Cell signalling 7076P2), HRP-conjugated anti-rabbit IgG (1:5000, Santa Cruz Biotechnology), Mouse α-β-actin (abcam ab8226).

## DATA AVAILABILITY

There are no restrictions on data availability. The RNA and DNA sequencing data presented in this study have been deposited at GEOs: GSEXXXXX.

## ACKNOWLEDGEMENTS.

We thank A. Hammarberg, S. da Rocha Baez, E. Monni, M. Persson-Vejgården, J. Nelander and S. Tesson for their technical assistance, S. Henikoff for the gift of the pA-MNase protein used in initial hNPC CUT&RUN experiments and the pAG-MNase plasmid (Addgene #123461) for in house production of the pAG-MNase used in all subsequent experiments. We thank P. Lehner for providing lentiviral vectors to knockout MORC2 and re-express MORC2 variants, the SCC Cell and Gene Therapy Core at Lund University and iPSC core at Karolinska Institute for generation of MORC2 edited iPSC clones. We acknowledge Clinical Genomics Lund and the Center for Translational Genomics (CTG) at Lund University for providing expertise and service with sequencing and analysis. This work was supported by the Swedish Government Initiative for Strategic Research Areas (MultiPark & StemTherapy) and grants from the Swedish Research Council (VR, 2022-02673 to J.J., 2021-03494 to C.H.D.), the Swedish Brain Foundation (Hjärnfonden, FO2023-0232, FO2023-0229 to C.H.D.), the Swedish Society for Medical Research (SSMF, S19-0100 to C.H.D.), Cancerfonden (222185Pj to J.J.), Barncancerfonden (PR2023-0099 to J.J.) and the Wellcome Trust (101908/Z/13/Z and 217191/Z/19/Z to Y.M.). This project has been made possible in part by grant 2023-331773 to C.H.D. from the Chan Zuckerberg Initiative DAF, an advised fund of the Silicon Valley Community Foundation. Finally, we also acknowledge The Royal Physiographic Society of Lund (project grant to F.D.), the Crafoord Foundation and Bente-Rexed-Gerstedt Foundation (project grants to C.H.D.).

## AUTHOR CONTRIBUTIONS

AA made and validated the SHSY5Y MORC2 KO cell line and complemented cell lines. FD and LC-V designed and performed iPSC culture and cerebral organoid differentiations. FD, SK, SB and AK performed CUT&RUN experiments. FD, LC-V and CD-H performed quality control analysis of iPSC lines. FD and JM conducted L1 CRISPRi experiments, validations and 2D NPC differentiations, with expert advice from AAd. CF prepared DNA for Nanopore sequencing and performed bioinformatic analysis of the data. CF, NP and SK processed Illumina sequencing experiments and performed bioinformatic analysis of the data. OEK, CD-H and JM tested 2D neural differentiation protocols. FD, LC-V and AAd performed nuclear isolation of organoids. JGJ prepared all Illumina sequencing libraries for bulk and single-nuclei RNA-seq and CUT&RUN experiments. CHD conceived and designed the study. YM, JJ and CHD supervised the study and acquired funding. FD, CF and CHD wrote the manuscript and made figures with assistance from NP. All authors reviewed the final version.

## SUPPLEMENTARY FIGURES

**Supplementary Figure 1.**
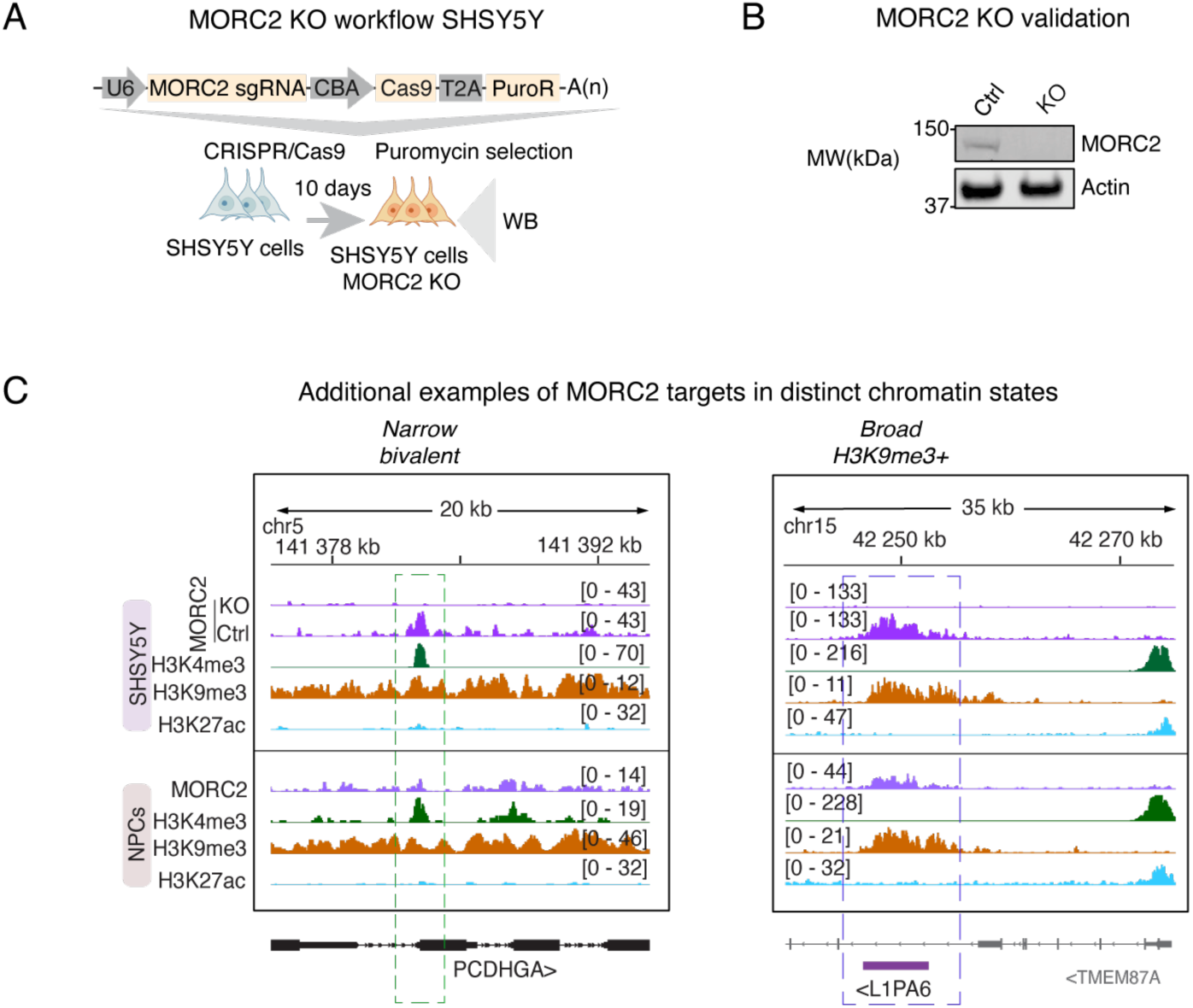
**(A)** Workflow for generating the MORC2 KO SHSY5Y cell line. **(B)** Validation of MORC2 KO by Western blot. **(C)** Further examples of the different chromatin states characterizing MORC2 targets, see also Fig 1.

**Supplementary Figure 2.**
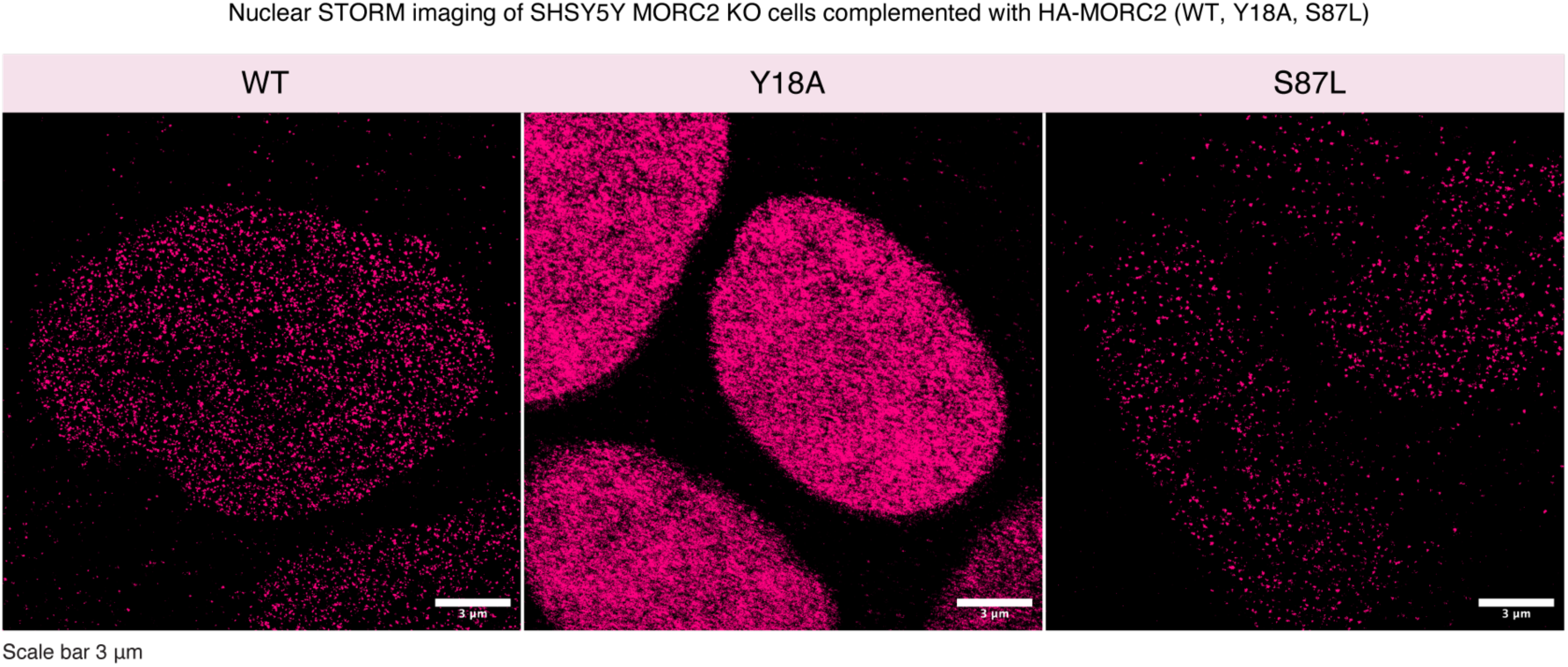
Nuclear STORM imaging of SHSY5Y MORC2 KO cells complemented with HA-MORC2 (WT, Y18A, S87L) targeting the HA epitope tag.

**Supplementary Figure 3.**
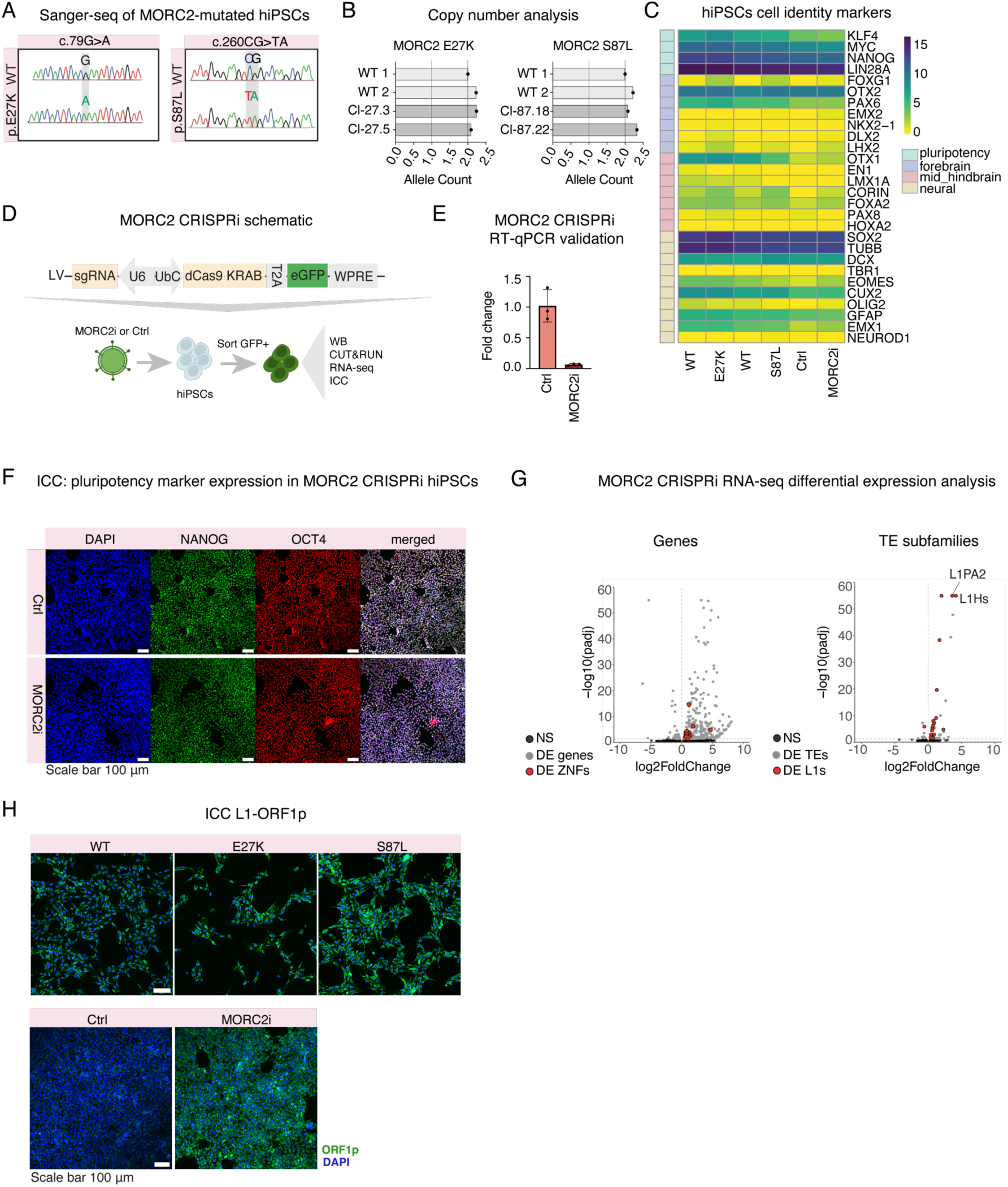
**(A)** Electropherogram of the Sanger sequencing confirming the CRISPR-directed MORC2 mutations (E27K and S87L) in representative clones **(B)** Copy number analysis showing allele count for each MORC2 mutant clone and WT hiPSC lines. **(C)** Unscaled heatmap showing normalized read counts of pluripotency and neural marker genes as calculated by DESeq2 in the hiPSCs used in the study. Two batches of RNA from each clone was used for WT (n=4), E27K (n=4) and S87L (n=4) genotypes. **(D)** Schematic workflow of MORC2 CRISPRi experiment in hiPSCs and subsequent analyses. **(E)** Validation of CRISPRi through RT-qPCR analysis of MORC2 expression in control (n=3) and CRISPRi (n=3) hiPSCs. **(F)** Immunohistochemistry of pluripotency markers (NANOG, OCT4) for MORC2 control and CRISPRi cell lines. **(G)** Volcano plot showing the results of the differential expression analysis (DESeq2) for genes (left) and TE subfamilies using TEtranscripts (right) in the MORC2 CRISPRi cell line (n=3) compared to control (n=3). Differentially expressed genes and LINE1 subfamilies are highlighted in red, other differentially expressed genes and subfamilies are presented in grey (padj <0.05). **(H)** Immunohistochemistry of L1-ORF1p (green) in E27K and S87L MORC2 mutant and CRISPRi cell lines with their respective controls. Experiments were performed at least twice with similar results.

**Supplementary Figure 4.**
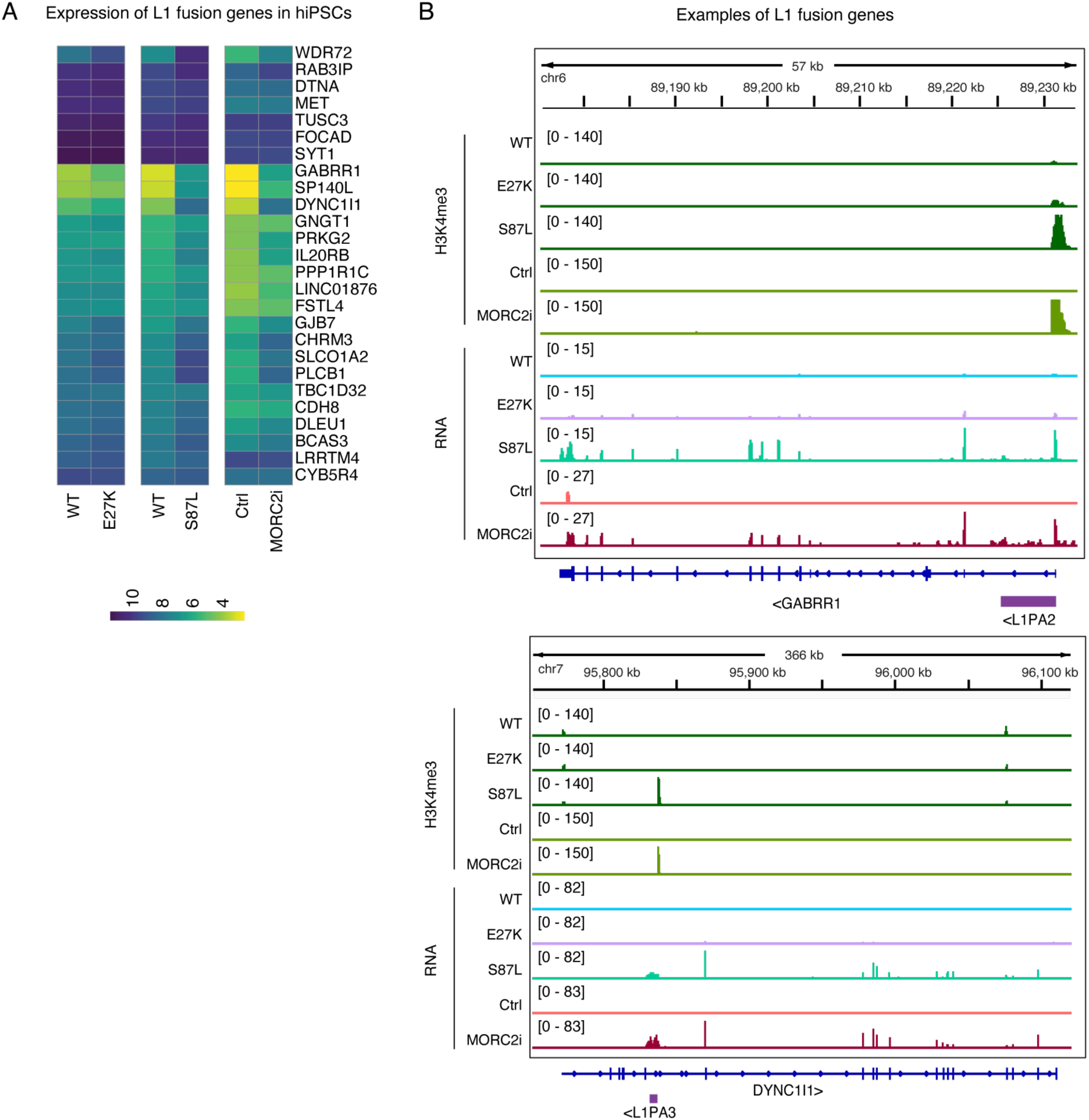
**(A)** Unscaled heatmap showing the normalized read counts of L1 fusion genes in the E27K (n=4) and S87L (n=4) MORC2 mutant iPSC lines as compared to WT controls (n=4) and MORC2 CRISPRi (n=3) iPSCs compared to its control (n=3). **(B)** Genome browser snapshots showing two examples of L1 fusion genes derepressed in MORC2 mutant and CRISPRi treatments.

**Supplementary Figure 5.**
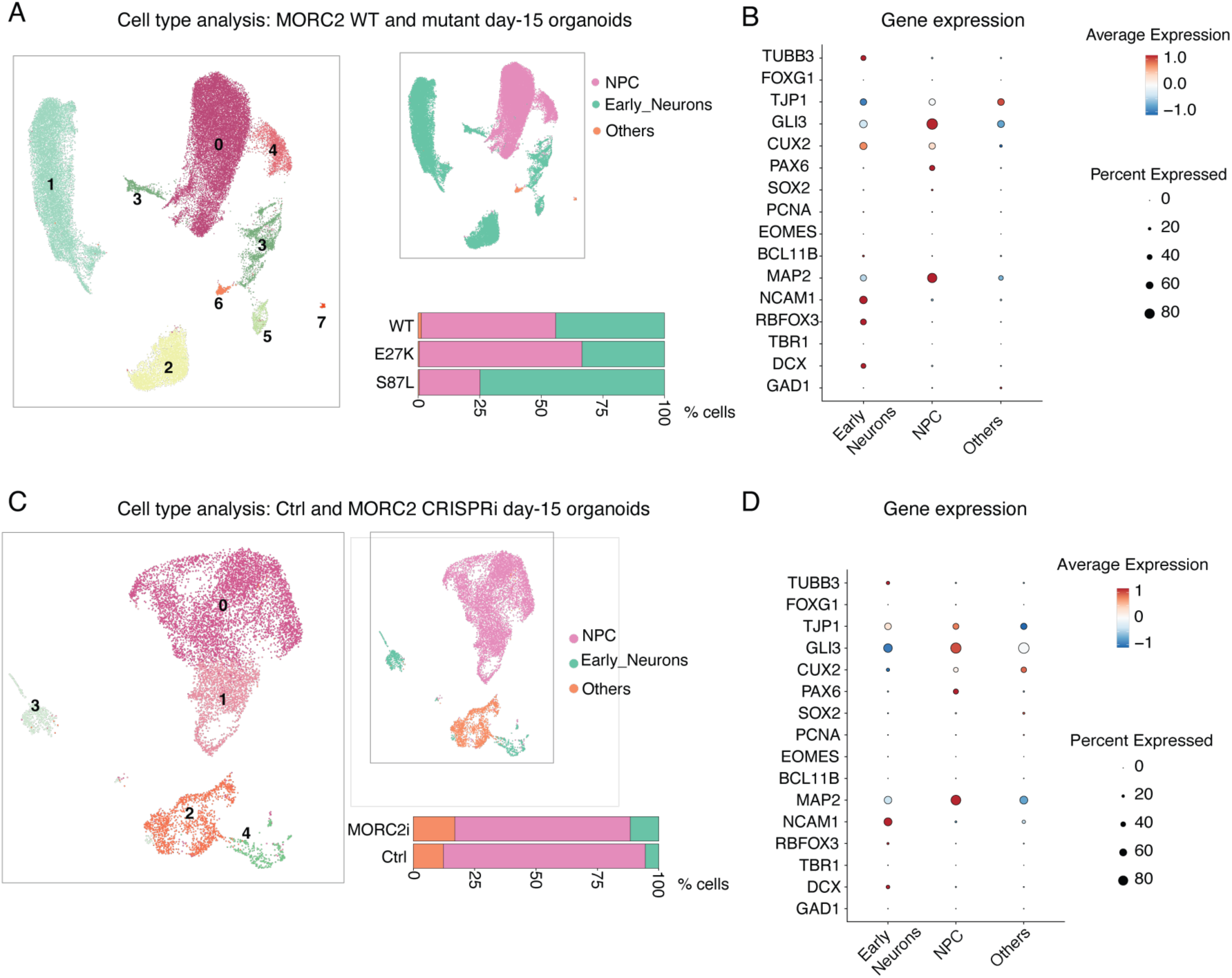
Cell composition of day 15, unguided cerebral organoids. **(A)** Left/Middle: UMAP showing the cell types identified in MORC2-WT (4-5 organoids from n=4 batches/clones), MORC2-E27K (4-5 organoids from n=2 batches/clones) and MORC2-S87L mutant (4-5 organoids from n=2 batches/clones) unguided cerebral organoids. Bottom: bar plot showing percentage of the cell type composition of the MORC2 WT and MORC2-S87L mutant cerebral organoids. **(B)** Dot plot showing neuronal and NPC gene markers used to characterize the cell clusters. Dot size corresponds to the proportion of cells expressing the gene. Dot color indicates average expression in each cell type. **(C)** Left/Middle: UMAP showing the cell types identified in control (4-5 organoids from n=1 batch) and MORC2 CRISPRi (4-5 organoids from n=1 batch) unguided cerebral organoids. Bottom: bar plot showing percentage of the cell type composition of the control and MORC2 CRISPRi cerebral organoids. **(D)** Dot plot showing neuronal and NPC gene markers used to characterize the cell clusters. Dot size corresponds to the proportion of cells expressing the gene. Dot color indicates average expression in each cell type.

**Supplementary Figure 6.**
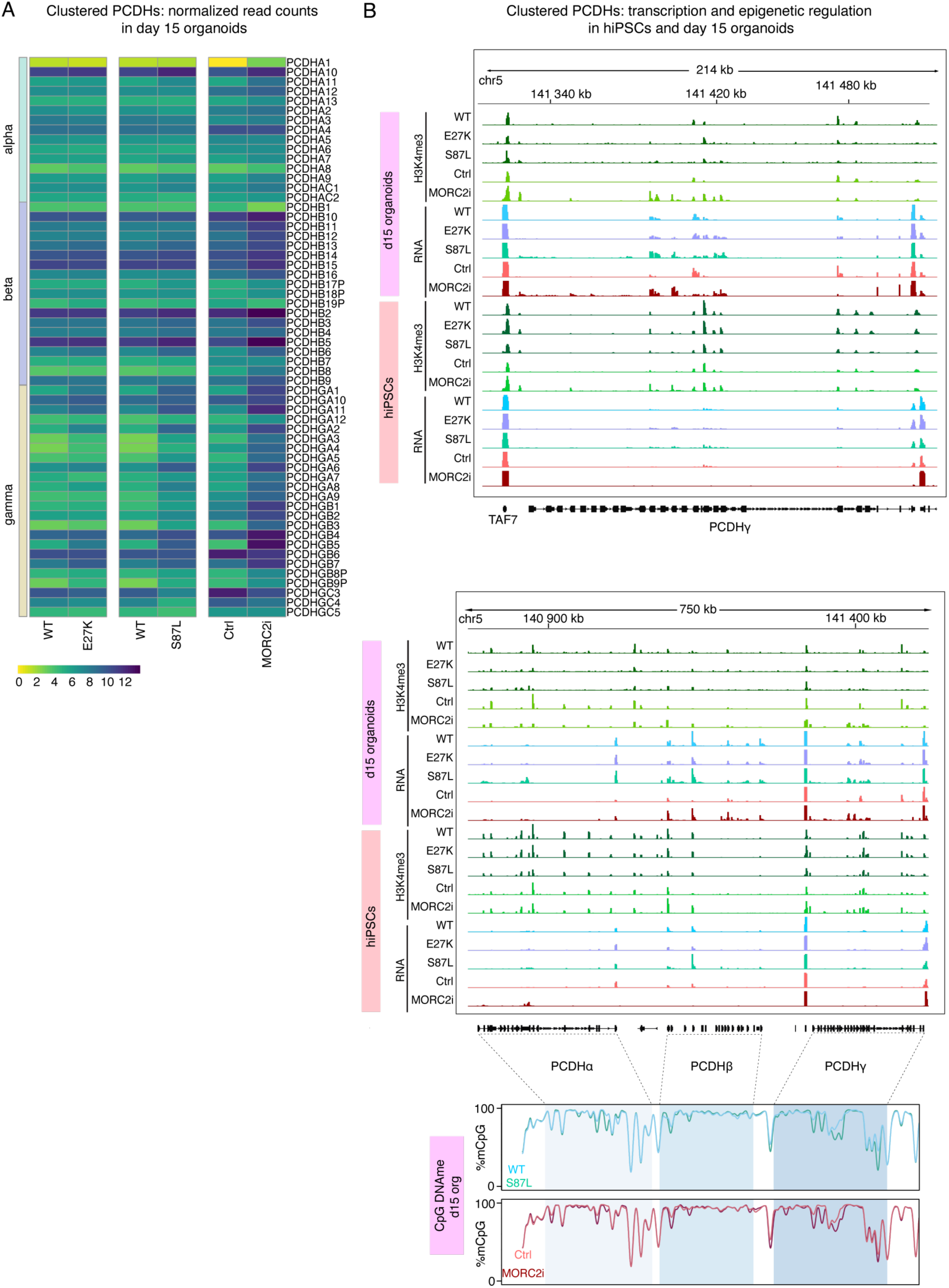
**(A)** Unscaled heatmap showing normalized read counts of bulk RNA-seq data for clustered protocadherin genes (PCDHα, PCDHβ and PCDHγ) as calculated by DESeq2 in WT (n=9 samples from a total of 27 organoids differentiated from 3 iPSC lines across 3 batches,) E27K (n=9 samples from a total of 27 organoids differentiated from 2 iPSC lines across 2 batches) and S87L (n=9 samples from a total of 27 organoids differentiated from 2 iPSC lines across 2 batches) cell lines, and from Control (n=3 samples derived from 9 organoids differentiated from 1 bulk transduced cell line across 1 batch) and MORC2 CRISPRi (n=3 samples derived from 9 organoids differentiated from 1 bulk transduced cell line across 1 batch). **(B)** Genome browser view of bulk RNA-seq and H3K4me3 CUT&RUN from organoid experiments of the PCDHγ cluster (top) and PCDHα, PCDHβ and PCDHγ clusters (center), and their mean percentage methylation profile (bottom). Data are representative of all replicates performed.

**Supplementary Figure 7.**
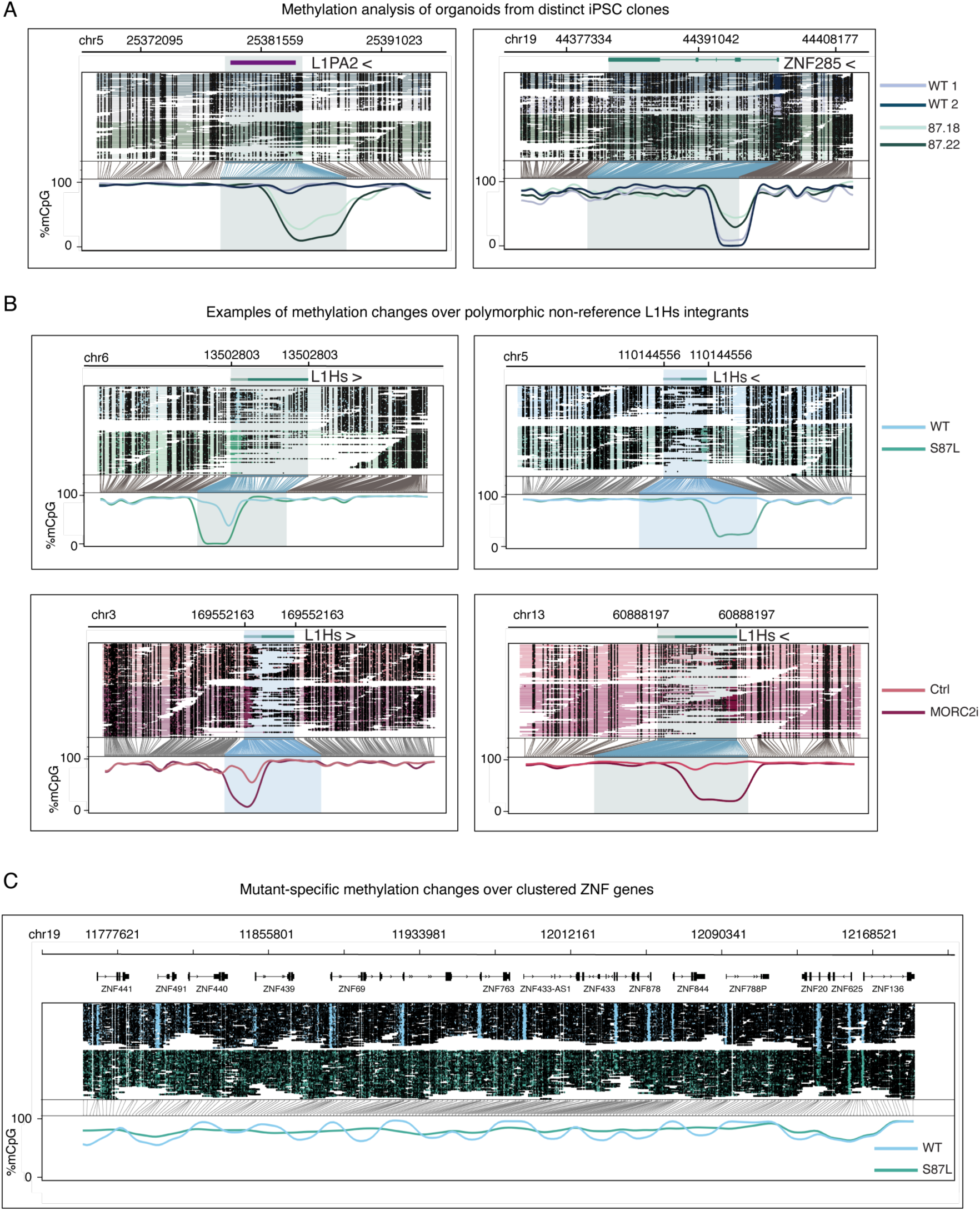
**(A)** Aligned Nanopore reads showing CpG methylation across clones belonging to the WT (n=2) and S87L (n=2) cell lines over example L1 and ZNF gene loci. **(B)** Aligned Nanopore reads showing CpG methylation over examples of non-reference LINE1 insertions in WT and S87L cell lines, and Ctrl and MORC2 CRISPRi cell lines. **(C)** Aligned Nanopore reads showing long-range dysregulated CpG methylation over a ZNF gene cluster.

**Supplementary Figure 8.**
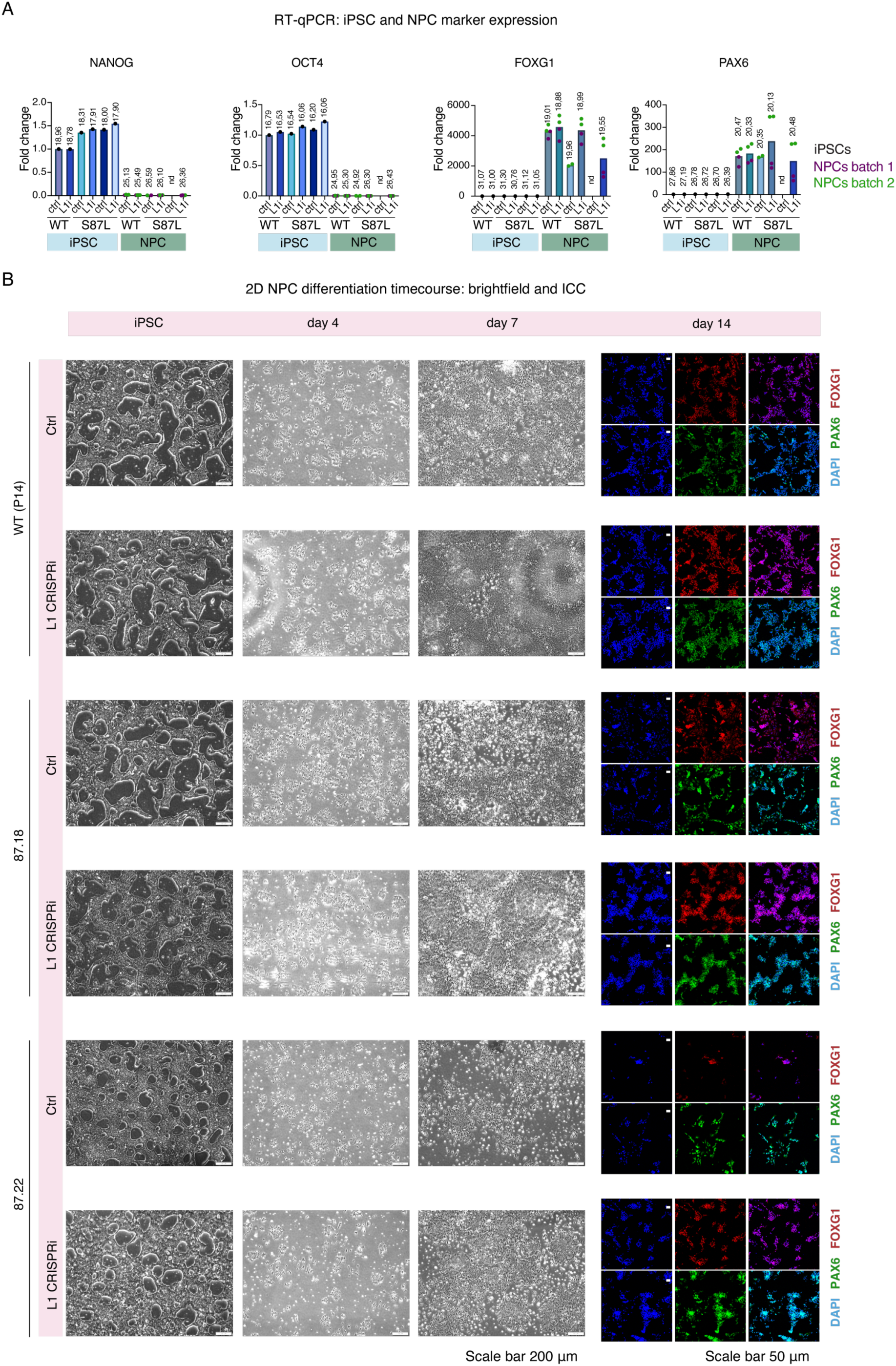
**(A)** RT-qPCR data from indicated iPSC (n=1) and NPCs derived therefrom (n=2 from two separate differentiation batches) to assess expression of pluripotency and neural progenitor markers. Each value is the mean of three technical replicates. nd, not determined due to insufficient RNA yield from the 87.22-Ctrl condition. In all cases Ct values were normalized to the *ACTB* housekeeping gene and fold-change calculated relative to the WT-Ctrl condition. **(B)** Representative brightfield and immunocytochemistry analysis of NPC differentiations from six cell lines at the indicated timepoints. Note, data shown for the WT-Ctrl line is the same as that shown in figure 6.

**Supplementary Figure 9.**
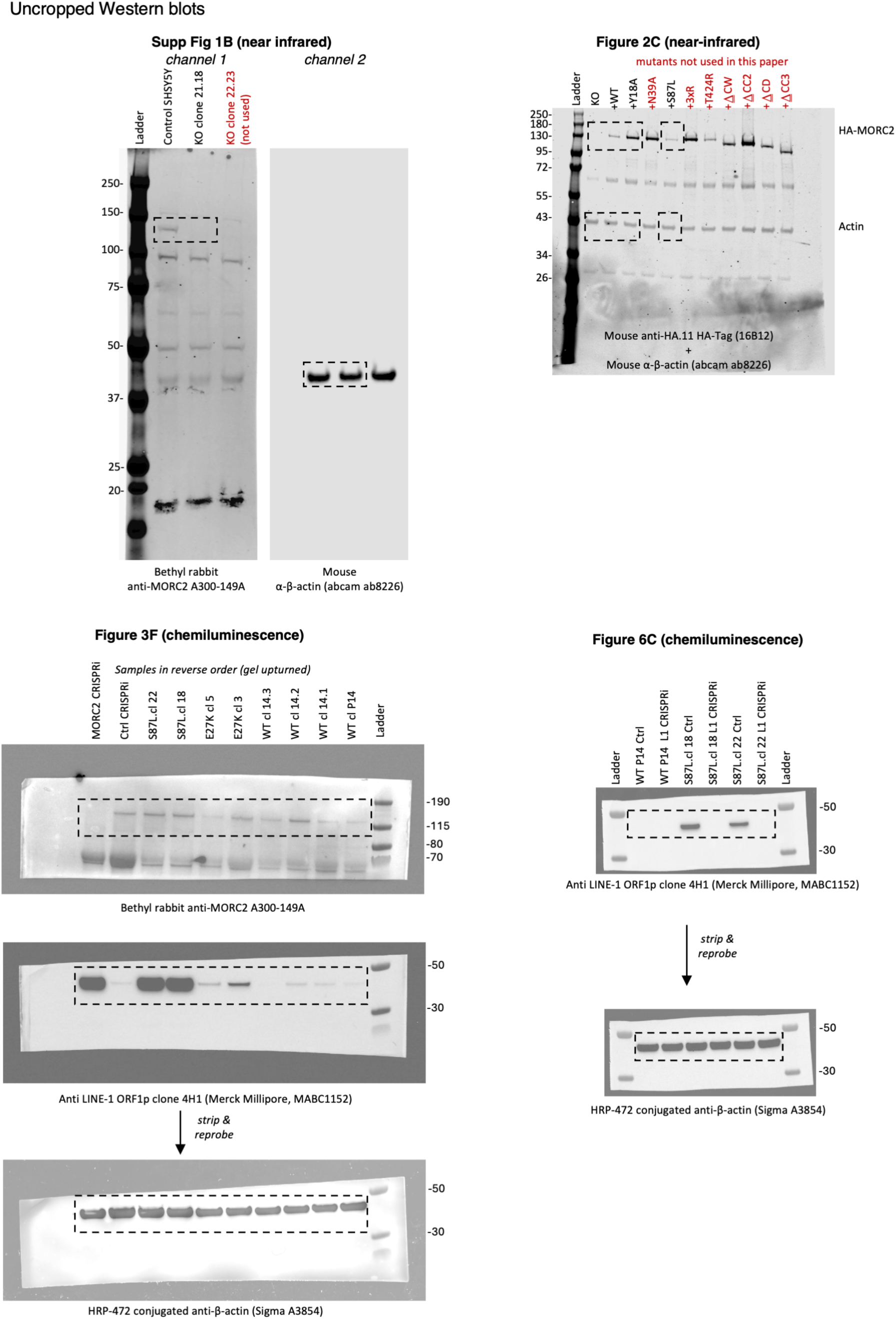
Uncropped Western blots used in the given figure panels. Where samples were not used in the study the labels are written in red. See also Methods.

